# Methyl ketone production by *Pseudomonas putida* is enhanced by plant-derived amino acids

**DOI:** 10.1101/496497

**Authors:** Jie Dong, Yan Chen, Veronica Teixeira Benites, Edward E.K. Baidoo, Christopher J. Petzold, Harry R. Beller, Aymerick Eudes, Henrik V. Scheller, Paul D. Adams, Aindrila Mukhopadhyay, Blake A. Simmons, Steven W. Singer

**Affiliations:** Joint BioEnergy Institute, Emeryville, CA, 94608; Biological Systems and Engineering Division, Lawrence Berkeley National Laboratory, Berkeley, CA, 94720; Earth and Environmental Sciences Area, Lawrence Berkeley National Laboratory, Berkeley, CA, 94720; Environmental Genomics and Systems Biology Division, Lawrence Berkeley National Laboratory, Berkeley, CA, 94720; Molecular Biophysics and Integrated Bioimaging Division, Lawrence Berkeley National Laboratory, Berkeley, CA, 94720

**Keywords:** lignin-related aromatics, methyl ketones, biomass hydrolysates, protein, amino acids

## Abstract

Plant biomass is an attractive source of renewable carbon for conversion to biofuels and bio-based chemicals. Conversion strategies often use a fraction of the total biomass, focusing on sugars from cellulose and hemicellulose. Strategies that use plant components such as plant-derived aromatics and amino acids have the potential to improve the efficiency of overall biomass conversion. *Pseudomonas putida* is a promising host for biomass conversion for its ability to metabolize a wide variety of organic compounds, including aromatics derived from lignin. *P. putida* was engineered to produce medium chain methyl ketones, which are promising diesel blendstocks and potential platform chemicals, from glucose and lignin-related aromatics, 4-hydroxybenzoate (4-HB) and protocatechuate (PCA). Unexpectedly, *P. putida* methyl ketone production was enhanced 2-to 5-fold compared to sugar controls when *Arabidopsis thaliana* hydrolysates derived from engineered plants that overproduce 4-HB and PCA, while *E. coli* production was lowered in these hydrolysates. This enhancement was more pronounced (~7-fold increase) with hydrolysates derived from non-engineered switchgrass (*Panicum virgatum* L.) suggesting it did not arise from overproduction of 4-HB and PCA. Global proteomic analysis of the methyl ketone-producing *P. putida* suggested that plant-derived amino acids may be the source of this enhancement. Mass spectrometry-based measurements of plant-derived amino acids demonstrated a high correlation between methyl ketone production and amino acid concentration in plant hydrolysates. Amendment of glucose-containing minimal media with a defined mixture of amino acids similar to those found in the hydrolysates studied led to a 9-fold increase in methyl ketone titer (1.1 g/L).

## INTRODUCTION

Plant biomass is an abundant potential resource for the production of biofuels and bio-based chemicals (Fiorentino, Ripa, & Ulgiati, 2017). The secondary plant cell walls are mainly composed of polysaccharides (cellulose, hemicellulose) and lignin, a complex polymer synthesized from aromatic monomers (Pandey & Kim, 2011). Conversion and upgrading of plant biomass has focused on C6 (cellulose) and C5 (hemicellulose) sugars obtained by physiochemical pretreatment and enzymatic hydrolysis of the cell-wall polysaccharides. The residual lignin is often combusted for its heating value; however, multiple methods exist to depolymerize lignin by chemical and biological processes (Zakzeski, Bruijnincx, Jongerius, & Weckhuysen, 2010)(Bugg, Ahmad, Hardiman, & Singh, 2011). These depolymerized lignins can be converted by microorganisms that can use these monoaromatic lignin-related compounds as bioconversion substrates (Beckham, Johnson, Karp, Salvachúa, & Vardon, 2016). Recent work has taken advantage of the ability of certain soil bacteria, particularly *Pseudomonas putida* and *Rhodococcus opacus*, to funnel lignin-related monoaromatics towards the production of polyhydroxyalkanoates, triacylglycerides and *cis*, *cis*-muconic acid (Linger et al., 2014)(Andreoni, Bernasconi, Bestetti, & Villa, 1991)(Vardon et al., 2015).

Bioconversion studies with lignin-related aromatics have focused on purified substrates (*p*-coumarate, 4-hydroxybenzoate, vanillate) or mixtures of aromatics obtained by chemical depolymerization of lignin. *Arabidopsis thaliana* and tobacco plants have been engineered with bacterial enzymes that shunt intermediates of lignin biosynthesis to alter lignin structures and lower plant lignin content (Eudes et al., 2015)(Eudes et al., 2012)(Wu et al., 2017). These engineered plants display an increase in saccharification efficiency when treated with cellulase/xylanase mixtures. As a byproduct of these transformations, soluble monoaromatics, which were present in the plant as aromatic glucosides, are produced at 1-5% of the total biomass. These soluble aromatics were extracted with organic solvent and treated under mild acidic conditions to release the deglycosylated monoaromatics (Eudes et al., 2015). Monoaromatics extracted from tobacco plants expressing dehydroshikimate dehydratase (QsuB) from *Corynebacterium glutamicum* were highly enriched in PCA(~5% of total biomass), and these extracts were incubated with engineered *Escherichia coli* strains that produced *cis, cis*-muconic acid (Wu et al., 2017). Therefore, these engineered plants may provide a direct method to produce lignin-related monoaromatics without requiring energy-intensive thermochemical treatments of lignin in order to liberate them.

Products obtained by engineering microbial fatty acid biosynthetic pathways (C_10_-C_18_) are attractive targets as precursors to biofuels and biochemicals. Free fatty acids, which have been overproduced in multiple microbes, can be converted to hydrocarbons and fatty acid ethyl esters, which are useful as diesel replacements or blendstocks (Beller, Lee, & Katz, 2015). Fatty alcohols, which are derived from free fatty acids by reduction, can be used as surfactants and lubricants (Espaux et al., 2017). Decarboxylation of beta-keto acids, an intermediate in fatty acid beta-oxidation, produces methyl ketones, which can also serve as diesel blendstocks as well as ingredients in the flavor and fragrance industry (Goh, Baidoo, Keasling, & Beller, 2012). Methyl ketones are particularly attractive as biofuel targets because they freely diffuse from cells and can be captured in an organic solvent overlay. Unlike free fatty acids, methyl ketones do not require additional processing steps to be blended into conventional diesel fuels. Methyl ketone production has been most intensively studied in *E. coli*, for which a titer of 3.4-5.4 g/L was achieved by fed-batch glucose fermentation (Goh et al., 2014)(Goh, Chen, Petzold, Keasling, & Beller, 2018). Methyl ketone production has also been demonstrated under fed-batch glucose fermentation in oleaginous yeast *Yarrowia lipolytica* (315 mg/L) (Hanko et al., 2018) and under autotrophic conditions from H_2_/CO_2_ in *Ralstonia eutropha* (180 mg/L) (Müller et al., 2013).

Here we describe the engineering of *P. putida* to produce medium chain methyl ketones from both glucose and lignin-related aromatics. Unexpectedly, methyl ketone production experiments with hydrolysates obtained from engineered *A. thaliana* plants that overproduce monoaromatics demonstrated significant improvements in methyl ketone production. These improvements were correlated with amino acid levels in the hydrolysate, rather than the presence of increased levels of these monoaromatics.

## MATERIALS AND METHODS

### Bacterial strains, media, and cultivation

*Pseudomonas putida* mt-2 (ATCC 33015) and *E. coli* S17-1 (ATCC 47055) were purchased from ATCC. *E. coli* DH5α was purchased from Thermo Fisher Scientific. *E. coli* EGS1895 (Goh et al., 2014) was obtained from the JBEI Registry (Table 1). *E. coli* S17-1 and *E. coli* DH5α were propagated at 37 °C in lysogeny broth (LB). Where necessary, medium was solidified with 1.0% (w/v) agar and supplemented with 50 μg/ml kanamycin. *P. putida* mt-2 and its engineered derivatives were grown at 30 °C in minimal medium (Rocha, da Silva, Taciro, & Pradella, 2008) (Ouyang, Liu, Fang, & Chen, 2007): (NH_4_)_2_SO_4_ 1.0 g/L, KH_2_PO_4_ 1.5 g/L, Na_2_HPO_4_ 3.54 g/L, MgSO_4_·7H_2_O 0.2 g/L, CaCl_2_·2H_2_O 0.01 g/L, ammonium ferric citrate 0.06 g/L and trace elements (H_3_BO_3_ 0.3 mg/L, CoCl_2_·6H_2_O 0.2 mg/L, ZnSO_4_·7H_2_O 0.1 mg/L, MnCl_2_·4H_2_O 0.03 mg/L, NaMoO_4_·2H_2_O 0.03 mg/L, NiCl_2_·6H_2_O 0.02 mg/L, CuSO_4_·5H_2_O 0.01 mg/L). Glucose, xylose, *p*-hydroxybenzoic acid or protocatechuic acid (pH was readjusted to 7.2) or biomass hydrolysate were supplemented as carbon source. *E. coli* EGS1895 was grown at 37 °C in M9-MOPS minimal medium (M9 medium supplemented with 75 mM MOPS, 2 mM MgSO_4_, 1 mg/L thiamine, 10 nM FeSO_4_, 0.1mM CaCl_2_, 56 mM NH_4_Cl_2_ and micronutrients including 3 mM (NH_4_)_6_Mo_7_O_24_, 0.4 mM boric acid, 30 mM CoCl_2_, 15 mM CuSO_4_, 80 mM MnCl_2_, and 10 mM ZnSO_4_) (Zhang, Carothers, & Keasling, 2012).

**Table 1.**
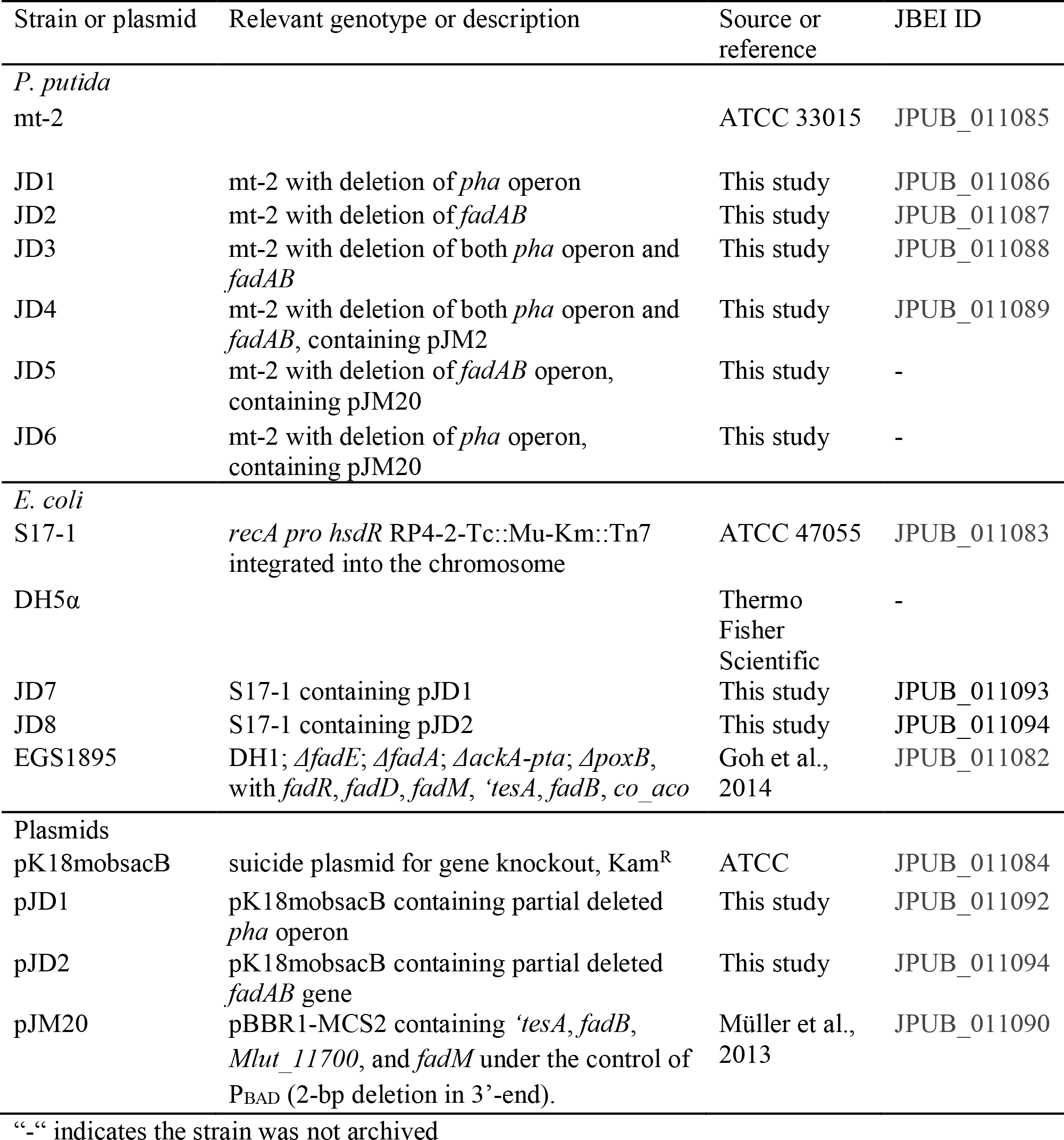
Bacterial strains and plasmids used in this study.

| Strain or plasmid | Relevant genotype or description | Source or reference | JBEI ID |
| --- | --- | --- | --- |
| *P. putida* |  |  |  |
| mt-2 |  | ATCC 33015 | JPUB_011085 |
| JD1 | mt-2 with deletion of *pha* operon | This study | JPUB_011086 |
| JD2 | mt-2 with deletion of *fadAB* | This study | JPUB_011087 |
| JD3 | mt-2 with deletion of both *pha* operon and *fadAB* | This study | JPUB_011088 |
| JD4 | mt-2 with deletion of both *pha* operon and *fadAB*, containing pJM2 | This study | JPUB_011089 |
| JD5 | mt-2 with deletion of *fadAB* operon, containing pJM20 | This study | - |
| JD6 | mt-2 with deletion of *pha* operon, containing pJM20 | This study | - |
| *E. coli* |  |  |  |
| S17-1 | *recA pro hsdR* RP4-2-Tc::Mu-Km::Tn7 integrated into the chromosome | ATCC 47055 | JPUB_011083 |
| DH5α |  | Thermo Fisher Scientific | - |
| JD7 | S17-1 containing pJD1 | This study | JPUB_011093 |
| JD8 | S17-1 containing pJD2 | This study | JPUB_011094 |
| EGS1895 | DH1; $\Delta fadE$; $\Delta fadA$; $\Delta ackA-pta$; $\Delta poxB$, with *fadR*, *fadD*, *fadM*, '*tesA*, *fadB*, *co_aco* | Goh et al., 2014 | JPUB_011082 |
| Plasmids |  |  |  |
| pK18mobsacB | suicide plasmid for gene knockout, Kam<sup>R</sup> | ATCC | JPUB_011084 |
| pJD1 | pK18mobsacB containing partial deleted *pha* operon | This study | JPUB_011092 |
| pJD2 | pK18mobsacB containing partial deleted *fadAB* gene | This study | JPUB_011094 |
| pJM20 | pBBR1-MCS2 containing '*tesA*, *fadB*, *Mlut_11700*, and *fadM* under the control of P<sub>BAD</sub> (2-bp deletion in 3'-end). | Müller et al., 2013 | JPUB_011090 |

“-“ indicates the strain was not archived

### Deletion of P. putida pha operon

The *pha* operon knockout of *P. putida* mt-2 was performed as previously described with some modifications (Ouyang, Liu, et al., 2007). *P. putida* mt-2 genome DNA was purified by QIAamp DSP DNA Mini Kit (Qiagen). A fragment from *P. putida* mt-2 genome containing a partial length of *phaC1*, and the whole length of *phaZ* and *phaC2* were amplified by PCR using the following two primers: 5’ primer AGAAAGCTTACCGGCAGCAAGGAC and 3’ primer GAGGCTAGCATCCAGTCAGCAGCTC. PCR products were digested by NheI and HindIII and then inserted in pK18mobsacB to form a new plasmid. The new plasmid was amplified in *E. coli* DH5α and completely digested by *PvuI*, and then the large fragment was self-ligated to form pJD1. As a result, partial 5’ sequence of *phaC1* (0.45 kbp) and partial 3’ sequence of *phaC2* (0.38 kbp) were inserted into pK18mobsacB in pJD1. Then pJD1 was transformed into *E. coli* S17-1 by electroporation. Transconjugations of *P. putida* mt-2 and *E. coli* S17-1 harboring recombinant plasmid pJD02 were carried out as previously described (Choi & Schweizer, 2005). Pseudomonas Isolation Agar (Difco) supplemented with 50 μg/mL kanamycin was used to select transconjugants. Then non-antibiotic LB agar plates supplemented with 250 g/L sucrose were used to select deletion mutants from transconjugants. The deletion mutants were verified by PCR using the same primers as those used in plasmid construction.

### Deletion of P. putida fadAB

The *fadAB* operon deletion in *P. putida* mt-2 was performed as described previously with some modifications (Ouyang, Luo, et al., 2007). A fragment from the *P. putida* mt-2 genome containing a partial length of *fadB* and *fadA* were amplified by PCR using the following two primers: 5’ primer ATTTCTAGAGCAGATGATGGCCTTC and 3’ primer CTGAAGCTTTGTAATGCCGGTATAC. PCR products and pK18mobsacB were double-digested by *XbaI* and *HindIII* and then ligated together to form a new plasmid. The new plasmid was completely digested by *SalI*, and then the large fragment was self-ligated to form pJD2. As a result, two DNA fragments, *fadB’* and *fadA'*, corresponding to a partial 5’ sequence of *fadB* and partial 3’ sequence of *fadA* was inserted into pK18mobsacB in pJD2. The homologous recombination of pJD04 into *P. putida* mt-2 chromosome and selection of knockout mutants were carried out as above in the *phaCZC* knockout. Repeating the above *fadAB* knockout procedure in the *phaCZC* deletion mutants provided a *P. putida* strain with both *fadAB* and *phaCZC* in-frame deletions.

### Transformation of methyl ketone production pathway into P. putida

The plasmid pJM20 encoding the methyl ketone production pathway was constructed previously (Müller et al., 2013). It contains the backbone from the broad-host-range vector pBBR1-MCS2 and has *‘tesA*, *fadB*, *Mlut_11700*, and *fadM* under the control of BAD promoter (P_BAD_). pJM20 was constructed by inserting a 2-bp deletion in the 3’-end of P_BAD_ to prevent inhibition of arabinose-induction in the presence of glucose (Miyada, Stoltzfus, & Wilcox, 1984). pJM20 was electroporated into *E.coli* DH5α for amplification (Johnson & Beckham, 2015) and was subsequently transferred into *P. putida* mt-2 mutants by electroporation (Johnson & Beckham, 2015). For transformation, 5 μL (0.2 −2 μg) of plasmid DNA was added to 50 μL of the electrocompetent cells, transferred to a chilled 0.2 cm electroporation cuvette, and electroporation was performed (1.6 kV, 25 uF, 200 Ω). SOC medium (450 μL, New England Biolabs, Ipswich, MA, USA) was added to the cells immediately after electroporation and the resuspended cells were incubated with shaking at 200 rpm, 30 °C for one hour. The entire transformation medium was plated on an LB agar plate containing 50 μg/mL kanamycin. Plasmid transformation was verified by restriction digest and gel electrophoresis.

### Methyl ketone production from purified substrates by P. putida and E. coli

Methyl ketone production assays were conducted in 15-mL test tubes with glucose, xylose, 4-hydroxybenzoate (4-HB) and protocatechuate (PCA) as carbon sources. Where indicated, the medium was amended with a mixture of amino acids (serine, valine, aspartate, phenylalanine, and tryptophan in equal amounts, total concentration was 0.5-1.5 g/L). A single colony of *P. putida* JD4 (*ΔfadAB*, *ΔphaCZC*, pJM20) was first grown at 30°C in minimal medium with 50 μg/mL kanamycin for ~12 h as the seed culture. The seed culture (5% v/v) was inoculated into each tube with 10 mL minimal medium. After ~6 h growth at 30 °C, 0.2 % (w/v) of L-arabinose was added for induction and 2 mL decane overlay was added for methyl ketone extraction. After 48 h, the decane was sampled and methyl ketone production measured using GC-MS (see below). *E. coli* EGS1895 freezer stock was first activated at 37 °C in M9-MOPS minimal medium with 50 μg/mL kanamycin for ~12 h as the seed culture. The seed culture (5% v/v) was inoculated into each tube with 10 mL minimal medium. After ~6 h growing at 37°C, 0.5 mM IPTG and 1 mM arabinose was added for induction and 2 mL decane overlay was added for methyl ketone extraction. After 72 h, the decane was sampled and methyl ketone production measured using GC-MS (see below).

### Preparation of biomass hydrolysates for methyl ketone production

*Arabidopsis thaliana* (L.) Heynh. ecotype Col-0 wild-type lines and engineered lines modified in lignin biosynthesis were previously described and grown at the Joint BioEnergy Institute (Eudes et al., 2012, 2015). Switchgrass (*Panicum virgatum* L., cultivar Alamo) was grown in a chamber at the Joint BioEnergy Institute under the following conditions: 25 °C, 60% humidity and 14 h of light per day (250 μmol.m^−2^.s^−1^). Sorghum (*Sorghum bicolor*) was provided by Idaho National Laboratory. Biomass (400 mg) was pretreated in 6.8 mL 1% (w/w) H_2_SO_4_ (6% w/w solid loading) at 121°C 20 psi for 1 h. Phosphate buffer (1 mL; pH 6.2) and distilled water (12 mL) were added and the pH was adjusted to 5.5 with an equimolar NaOH and KOH solution (5 N). CTec3 (5 μL, Novozymes) was added to the pH-adjusted slurry and saccharification was conducted in a VWR hybridization oven (Model 5420) at 15 rpm, 50 °C for 48 h. The pH was adjusted to 7.2 with equimolar NaOH and KOH solution (5 N) and the combined hydrolysate was centrifuged at 10,000 *x g* for 20 min and filtered through a 0.2-μm nylon membrane to remove residual solids. The clear supernatant was stored for 4 °C for further use. To separate the acid hydrolysate, the slurry obtained after dilute acid pretreatment was centrifuged at 10,000 *x g* for 20 min and the supernatant, referred to as the acid hydrolysate, stored at 4°C. The remaining solid was resuspended in 19 mL of distilled water and its pH adjusted to 5.5 with an equimolar NaOH and KOH solution (5 N). Hydrolysis with CTec3 was performed as described above, yielding the enzymatic hydrolysate. For methyl ketone production using these enzymatic hydrolysates, concentrated medium supplements were added into the hydrolysates, as the media lacked additional carbon sources and buffers (KH_2_PO_4_ and Na_2_HPO_4_ in *P. putida* medium; MOPS in *E. coli* medium). The solution was then filtered through a 0.2-μm membrane for sterilization and the cultivation performed under the same conditions as the pure substrates.

### Proteomics

Samples prepared for shotgun proteomic experiments were analyzed by an Agilent 6550 iFunnel Q-TOF mass spectrometer (Agilent Technologies, Santa Clara, CA) coupled to an Agilent 1290 UHPLC system as described previously (González Fernández-Niño et al., 2015). Peptides (20 μg) were separated on a Sigma–Aldrich Ascentis Peptides ES-C18 column (2.1 mm × 100 mm, 2.7 μm particle size, operated at 60°C) at a 0.40 mL/min flow rate and eluted with the following gradient: initial condition was 95% solvent A (0.1% formic acid) and 5% solvent B (99.9% acetonitrile, 0.1% formic acid). Solvent B was increased to 35% over 120 min, and then increased to 50% over 5 min, then up to 90% over 1 min, and held for 7 min at a flow rate of 0.6 mL/min, followed by a ramp back down to 5% B over 1 min where it was held for 6 min to re-equilibrate the column to original conditions. Peptides were introduced to the mass spectrometer from the liquid chromatography (LC) by using a Jet Stream source (Agilent Technologies) operating in positive-ion mode (3,500 V). Source parameters employed gas temp (250°C), drying gas (14 L/min), nebulizer (35 psig), sheath gas temp (250°C), sheath gas flow (11 L/min), VCap (3,500 V), fragmentor (180 V), OCT 1 RF Vpp (750 V). The data were acquired with Agilent MassHunter Workstation Software, LC/MS Data Acquisition B.06.01 operating in Auto MS/MS mode whereby the 20 most intense ions (charge states, +2–5) within 300–1,400 *m/z* mass range and above a threshold of 1,500 counts were selected for MS/MS analysis. MS/MS spectra (100– 1,700 *m/z*) were collected with the quadrupole set to “Medium” resolution and were acquired until 45,000 total counts were collected or for a maximum accumulation time of 333 ms. Former parent ions were excluded for 0.1 min following MS/MS acquisition. The acquired data were exported as mgf files and searched against the latest *P. putida* KT2440 protein database supplemented with coding protein sequences in the TOL plasmid with Mascot search engine version 2.3.02 (Matrix Science). The chromosomal genome of *P. putida* KT2440 and *P. putida* mt-2 are identical. The resulting search results were filtered and analyzed by Scaffold v 4.3.0 (Proteome Software Inc.). A total of 876 proteins were found that had at least two peptides identified with 95% confidence in at least one of the biological replicates. The normalized spectral counts of identified proteins were exported for relative quantitative analysis.

### Analysis

Concentrations of organic compounds except for amino acids in the media or hydrolysates were measured with an Agilent 1100 Series HPLC system equipped with an Agilent 1200 Series refractive index detector (RID) and diode array and multiple wavelength detector (DAD) (Agilent Technologies) (Goh et al., 2014). Aliquots (1 mL) of cell cultures were collected and centrifuged. The supernatants were filtered through a spin-cartridge with a 0.45-μm nylon membrane. For glucose and xylose detection, 5-μL samples were eluted through Aminex HPX-87H ion-exclusion column (300-mm length, 7.8-mm internal diameter; Bio-Rad Laboratories, Inc.) at 50 °C with 4 mM sulfuric acid at a flow rate of 600 μL/min for 15 min. Glucose and xylose were detected by RID. For 4-HB and PCA detection, 5 μL was eluted through Hypersil ODS C18 column (250-mm length, 4.6-mm internal diameter; Thermo Fisher Scientific) at 20°C with 20% (v/v) acetonitrile and 0.5% (v/v) acetic acid in water at a flow rate of 900 μL/min for 15 min. PCA and 4-HB were detected by DAD at 275 nm (Heinaru et al., 2001).

Methyl ketones present in the decane overlay were quantified using electron-ionization gas chromatography/mass spectrometry (GC/MS) as described previously (Goh et al., 2012). For the measurement of amino acids in hydrolysates, liquid chromatographic separation was conducted using a Kinetex HILIC column (100-mm length, 4.6-mm internal diameter, 2.6-μm particle size; Phenomenex, Torrance, CA) using a 1200 Series HPLC system (Agilent Technologies, Santa Clara, CA, USA). The injection volume for each measurement was 2 μL. The sample tray and column compartment were set to 6°C and 40°C, respectively. The mobile phase was composed of 20 mM ammonium acetate in water (solvent A) and 10 mM ammonium acetate in 90% acetonitrile and 10% water (solvent B) (HPLC grade, Honeywell Burdick & Jackson, CA, USA).

Ammonium acetate was prepared from a stock solution of 100 mM ammonium acetate and 0.7 % formic acid (98-100% chemical purity, from Sigma-Aldrich, St. Louis, MO, USA) in water. Amino acids were separated with the following gradient: 90% to 70%B in 4 min, held at 70%B for 1.5 min, 70% to 40%B in 0.5 min, held at 40%B for 2.5 min, 40% to 90%B in 0.5 min, held at 90%B for 2 min. The flow rate was varied as follows: held at 0.6 mL/min for 6.5 min, linearly increased from 0.6 mL/min to 1 mL/min in 0.5 min, and held at 1 mL/min for 4 min. The total run time was 11 min. The mass spectrometry parameters have been previously described (Bokinsky et al., 2013).

### Strain and Data Availability

Strains are available from the JBEI Public Registry (https://public-registry.jbei.org/) and the IDs are listed in Table 1. The mass spectrometry proteomics data have been deposited to the ProteomeXchange Consortium via the PRIDE partner repository (Vizcaíno et al., 2016) with the dataset identifier PXD012013 and 10.6019/PXD01201.

## RESULTS

### Engineering P. putida for methyl ketone production

A successful strategy for microbial production of methyl ketones (C_11_-C_15_) is to prevent native β-oxidation, deregulate fatty acid biosynthesis and express a truncated β-oxidation pathway to form β-keto-acids, which spontaneously decarboxylate to methyl ketones (Goh et al., 2012). For *P. putida*, previous work has demonstrated that the deletion of *fadAB* and genes for PHA polymerization (*phaC*) prevented β-oxidation of fatty acids and overproduced β-hydroxy-acids, which are thiolytic products of β-hydroxyacyl-CoAs and key intermediates in methyl ketone production (Ouyang, Luo, et al., 2007). Therefore, both *fadAB* and the PHA synthase *pha* operon (*phaC1*-*phaZ*-*phaC2*) were deleted to eliminate competition for the hydroxyacyl-CoA intermediates. Four genes (*‘tesA*, *Mlut_11700*, *fadB* and *fadM*) were transformed into *P. putida* mt-2 on a pBBR1-MCS2-based broad host plasmid under the control of P_BAD_ (Müller et al., 2013). *E. coli ‘tesA* encodes for a thioesterase with a truncated leader sequence that deregulates fatty acid biosynthesis by thiolysis of acyl-CoAs and acyl-ACPs. *Mlut_11700* encodes for a soluble acyl-CoA oxidase from *Micrococcus luteus*. *E. coli* FadB converts the enoyl-CoAs produced by the acyl-CoA oxidase to β-keto-acyl-CoAs, which are converted to β-keto acids by the *E. coli* thioesterase FadM. The spontaneous decarboxylation of β-keto acids produces methyl ketones through a non-enzymatic process (Goh et al., 2014).

### Methyl ketone production by P. putida

The production of methyl ketones was tested with either glucose or lignin-related aromatics as the sole carbon source in minimal medium. Two aromatics, 4-hydroxybenzoate (4-HB) and protocatechuate (PCA), were also tested as carbon sources since these aromatics accumulated in engineered *Arabidopsis* plants whose hydrolysates we planned on testing for methyl ketone production (Eudes et al., 2012)(Eudes et al., 2015). *P. putida* JD4, which contained the deletions in the PHA synthesis and β-oxidaton pathways, produced >200 mg/L of methyl ketones from glucose and ~300 mg/L from 4-HB. PCA was a poor substrate for methyl ketone production, producing <30 mg/L from the combined mutant, which may arise due to chelation of the Fe^2+^ present in the medium by PCA (Kamariotaki et al., 1994) (Gerega et al., 1987) (**Figure 1**). The chelation complex may hinder growth by reducing the availability of PCA or directly inhibiting *P. putida* and was therefore not tested in the other deletion strains. *P. putida* JD5, which has a deletion of *fadAB*, resulted in the production of 140-150 mg/L methyl ketones from glucose and 4-HB (**Figure S1**). *P. putida* JD6, which has the lesion in the PHA synthesis genes, only produced methyl ketones from 4-HB, which may arise from a reduced flux of aromatic substrates to PHA synthesis compared to glucose (Linger et al., 2014). The most abundant methyl ketones produced in *P. putida* were C_13_ and C_15_ methyl ketones, which is similar in chain length to *R. eutropha* (Müller et al., 2013) and *E. coli* (Goh et al., 2012). The majority of the methyl ketones (>60%) were unsaturated suggesting that they were converted from a high portion of unsaturated fatty acids produced in *P. putida*.

**Figure 1.**
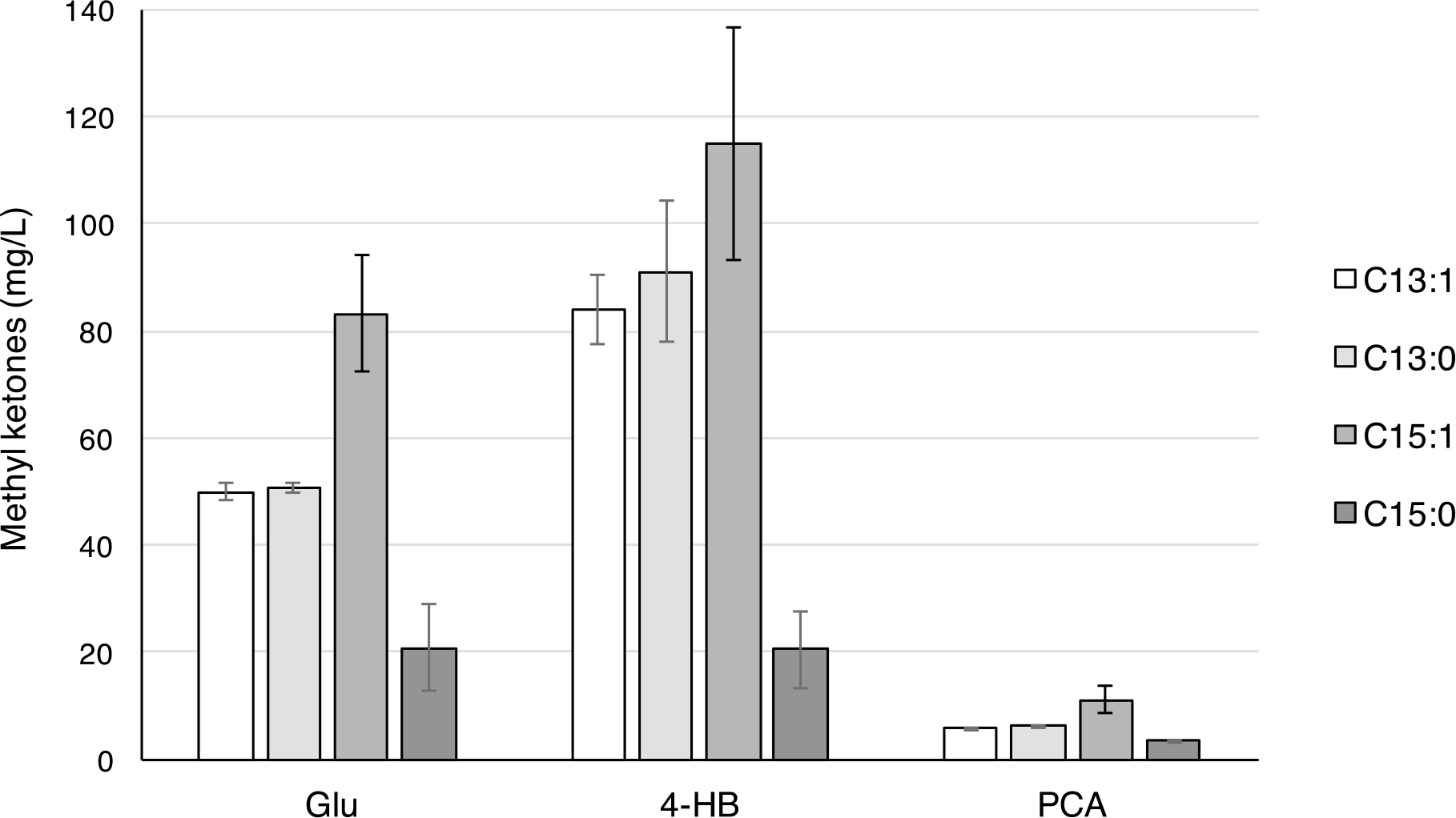
Methyl ketone production from *P. putida* strain JD4. *P. putida* JD4 was grown with 2% glucose (Glu), 1.5% *p*-hydroxybenzoate (4-HB) and 1% protocatechuate (PCA). Decane (2 mL) was overlaid onto the cultures when arabinose (0.2%) was added at 6 h to induce methyl ketone production. Titers are reported for methyl ketones in decane overlay at 48 h. Cultures were performed in triplicate and error reported as standard deviation.

### Methyl ketone production from engineered Arabidopsis thaliana

Methyl ketone production with single substrates provided background for experiments that tested production with plant hydrolysates. As noted above, we chose hydrolysates from *A. thaliana* lines in which lignin biosynthesis was disrupted by the expression of bacterial genes. Two strategies were pursued in *A. thaliana* to disrupt lignin biosynthesis, which lowered plant lignin content and improved polysaccharide hydrolysis. In the first strategy, expression of 3-dehydroshikimate dehydratase (QsuB from *C. glutamicum*) converts 3-dehydroshikimate, an important intermediate in lignin synthesis, into PCA (Eudes et al., 2015). In the second strategy hydroxycinnamoyl-CoA hydratase-lyase (HCHL) from *Pseudomonas fluorescens* AN103 was expressed to reroute the lignin biosynthetic intermediate *p*-coumaroyl-CoA into 4-hydroxybenzaldehyde, which was further oxidized to 4-HB (Eudes et al., 2012). The PCA-producing *A. thaliana* line is subsequently referred to as the QsuB line and the 4-HB-producing line is referred to as the HCHL line.

A mild acid pretreatment of biomass from these engineered plants followed by enzymatic hydrolysis was developed to produce hydrolysates that contained both sugars and aromatics. The two *A. thaliana* lines released more sugars, especially glucose, compared to wild type control plants, (**Figure 2A**). Hydrolysates from the QsuB line contained ~1.0 g/L of PCA (~5% of dry biomass) and those from HCHL line had ~ 0.2 g/L of 4-HB (~1% of dry biomass). Although the *A. thaliana* biomass also generates ~2.0 g/L xylose, primarily by acid hydrolysis, *P. putida* mt-2 cannot metabolize xylose into TCA cycle. Incubations with glucose and xylose demonstrated >90% conversion of xylose to xylonate, which was not further incorporated (data not shown). The near quantitative production of xylose from xylonate has previously been observed in *P. putida* S12 (Meijnen, de Winde, & Ruijssenaars, 2008).

**Figure 2.**
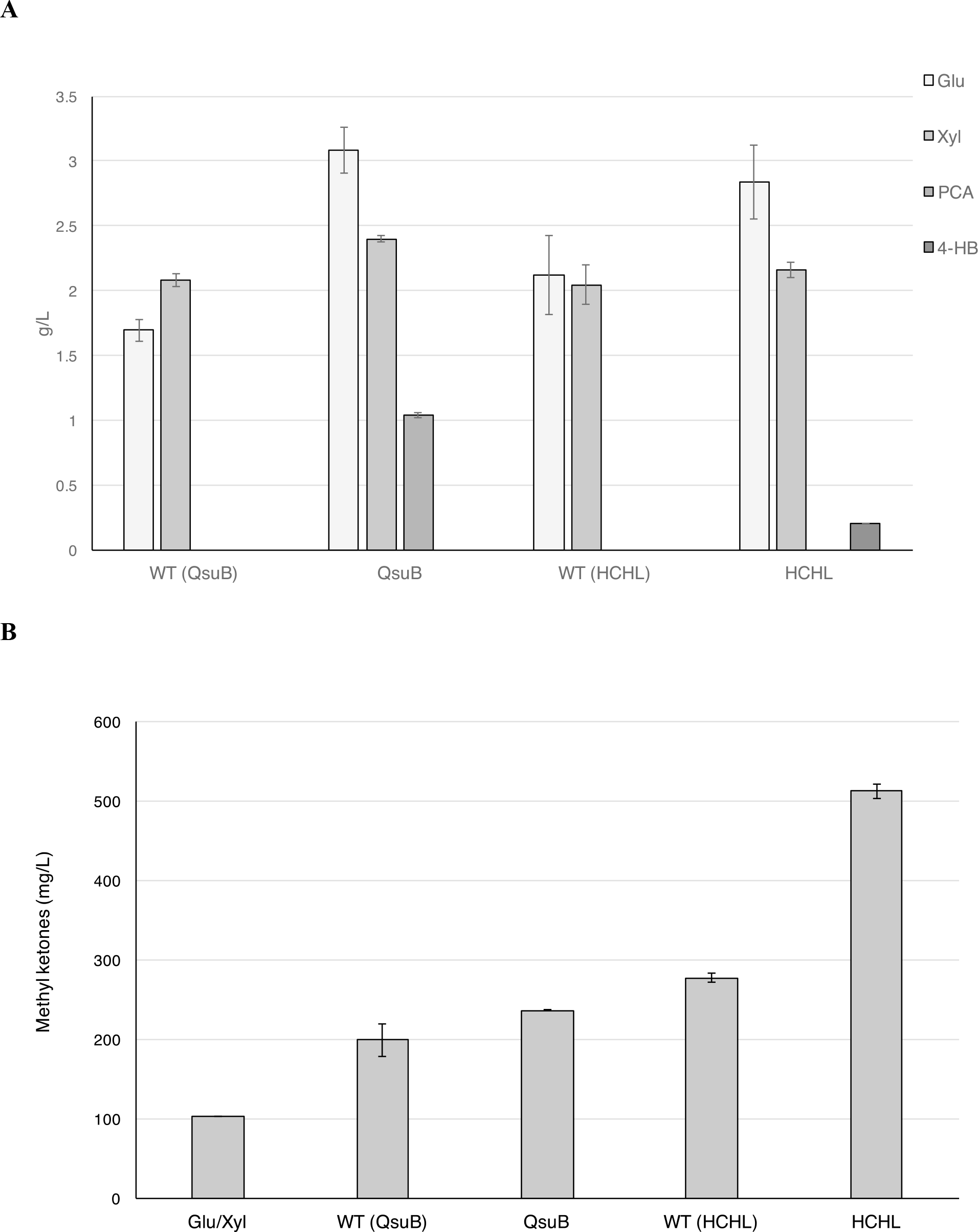
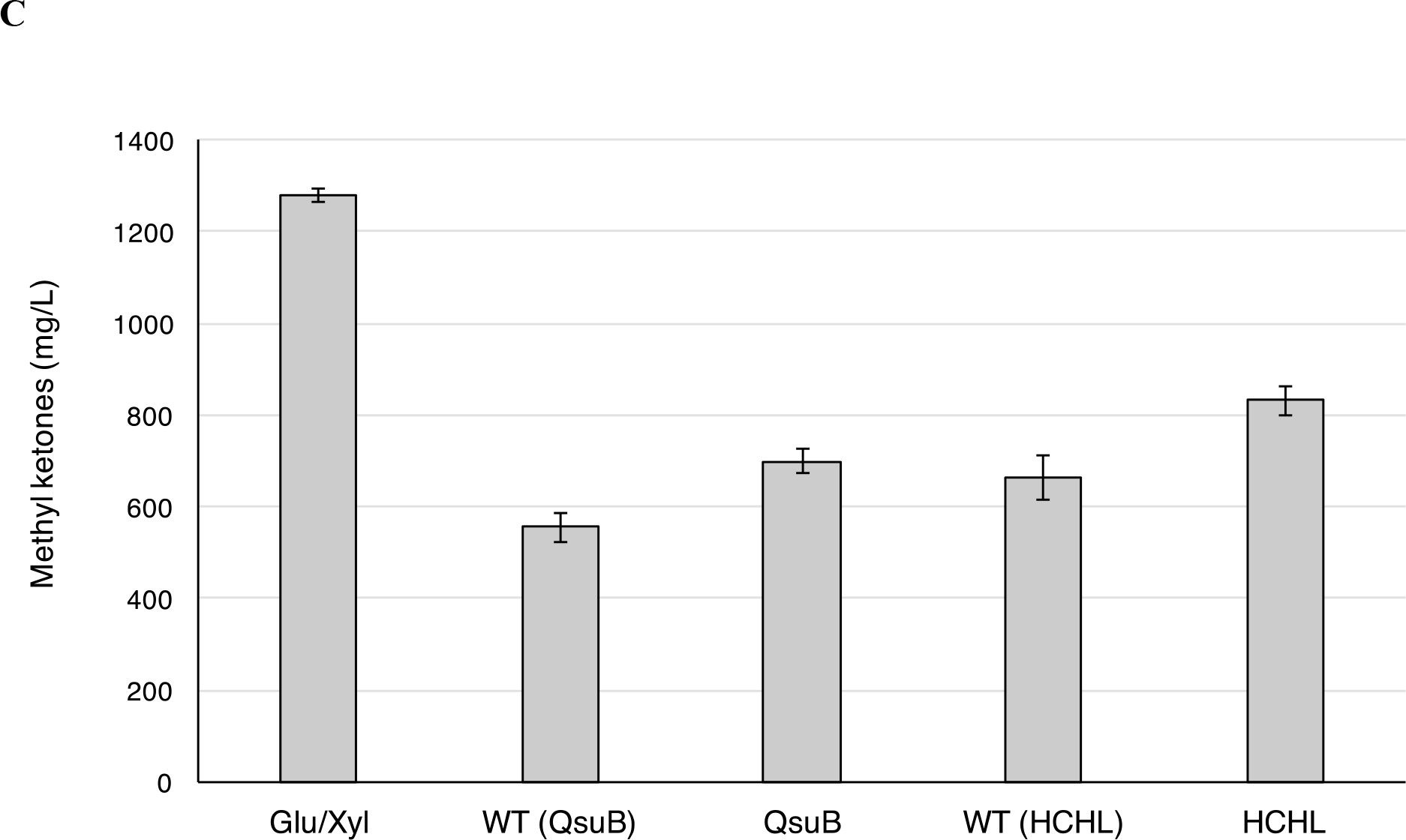
A) HPLC measurements of glucose (Glu), xylose (Xyl) and monoaromatics (4-HBA, PCA) released from *A. thaliana* biomass by sequential dilute acid and enzymatic hydrolysis. *A. thaliana* QsuB and HCHL lines were compared to their respective wild type (WT) controls. Measurements were performed in triplicate and error reported as ± standard deviation; B) Methyl ketone production by *P. putida* JD4 with hydrolysates derived from *A. thaliana* biomass. Cultures with *A. thaliana* hydrolysates were compared to a control containing 3 g/L of glucose and 2 g/L of xylose (Glu/Xyl). Methyl ketone production was performed as described in the Figure 1 legend; C) *E. coli* EGS1895 methyl ketone production with the same *A. thaliana* hydrolysates and Glu/Xyl control as the experiment with *P. putida* JD4.

The growth and methyl ketone production using these *A. thaliana* hydrolysates as carbon sources were tested. *P. putida* reached higher optical densities in the cultures grown with hydrolysates from the QsuB and HCHL lines compared to those grown with hydrolysates obtained from the corresponding wild type control plants, likely because of the increased concentration of glucose in these hydrolysates (**Figure S2A**). Surprisingly, all the *A. thaliana* hydrolysates produced higher levels of methyl ketones than the sugar-only controls (**Figure 2B**). Hydrolysates from wild type and QsuB plants provided 2-3 fold higher methyl ketone than the glucose/xylose control, which suggested that other components in the plant hydrolysates increased methyl ketone titers. The hydrolysate from the HCHL plants gave the highest methyl ketone titer (~ 500 mg/L), which is ~2x the titer achieved with hydrolysates from wild type plants and ~ 5x the titer of the sugar-only control.

Methyl ketone production with *P. putida* was compared to an *E. coli* strain (*E. coli* EGS1895) optimized for methyl ketone production from glucose by improving flux through the fatty acid pathway and eliminating acetate production; these modifications to the base strain, which focused on truncating β-oxidation, demonstrated >1g/L of methyl ketone production from 1% glucose in M9 minimal medium (Goh et al., 2014). As with the *P. putida* cultures, *E. coli* GS1895 reached higher optical density in the cultures grown with hydrolysates from the *A. thaliana* QsuB and HCHL lines (**Figure S2B**). *E. coli* EGS1895 displayed high levels of methyl ketone production (1.3 g/L) in cultures with the glucose/xylose control, but *E. coli*’s methyl ketone titers using *A. thaliana* hydrolysates were only 40%-60% of the control. As with *P. putida*, cultures grown on hydrolysates from biomass of the *A. thaliana* HCHL line produced the highest concentration of methyl ketones among the hydrolysate-fed cultures (~800 mg/L) (**Figure 2C**).

### Methyl ketones production from switchgrass

The attenuation of methyl ketone production by *E. coli* when grown on plant hydrolysates was not surprising based on previous studies demonstrating inhibition by plant hydrolysates; however, the large increase in titer compared to sugar only controls for *P. putida* was unexpected. To investigate further if the high methyl ketone production from hydrolysates was unique to *A. thaliana*, we tested hydrolysates obtained from switchgrass. Switchgrass has been shown to be a source of hydrolysates for biological conversion to a variety of biofuels and bio-based chemicals (Bokinsky et al., 2011). Mild dilute acid pretreatment and enzymatic hydrolysis yielded ~ 5.0 g/L of glucose and ~ 4.0 g/L of xylose in the combined hydrolysate (**Figure 3A**). In a second pretreatment, the solid was separated from the acid hydrolysate and subsequently hydrolyzed with enzymes, producing a separate enzymatic hydrolysate. The acid hydrolysate contained mainly xylose (~ 3.5 g/L) and small amount of glucose (~ 0.8 g/L), while glucose (~g/L) predominated in the enzymatic hydrolysate with a small amount of xylose (~ 0.8 g/L).

**Figure 3.**
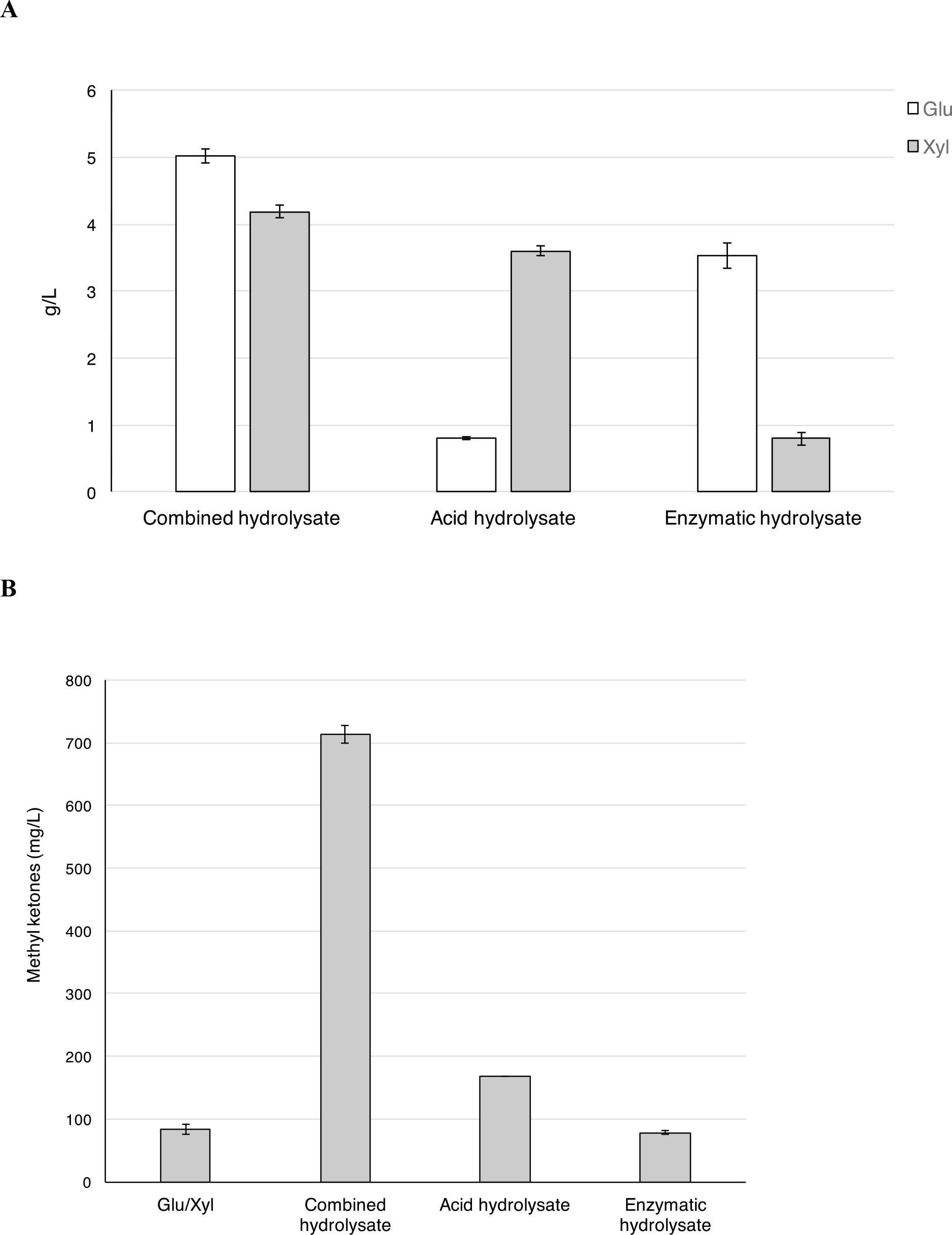
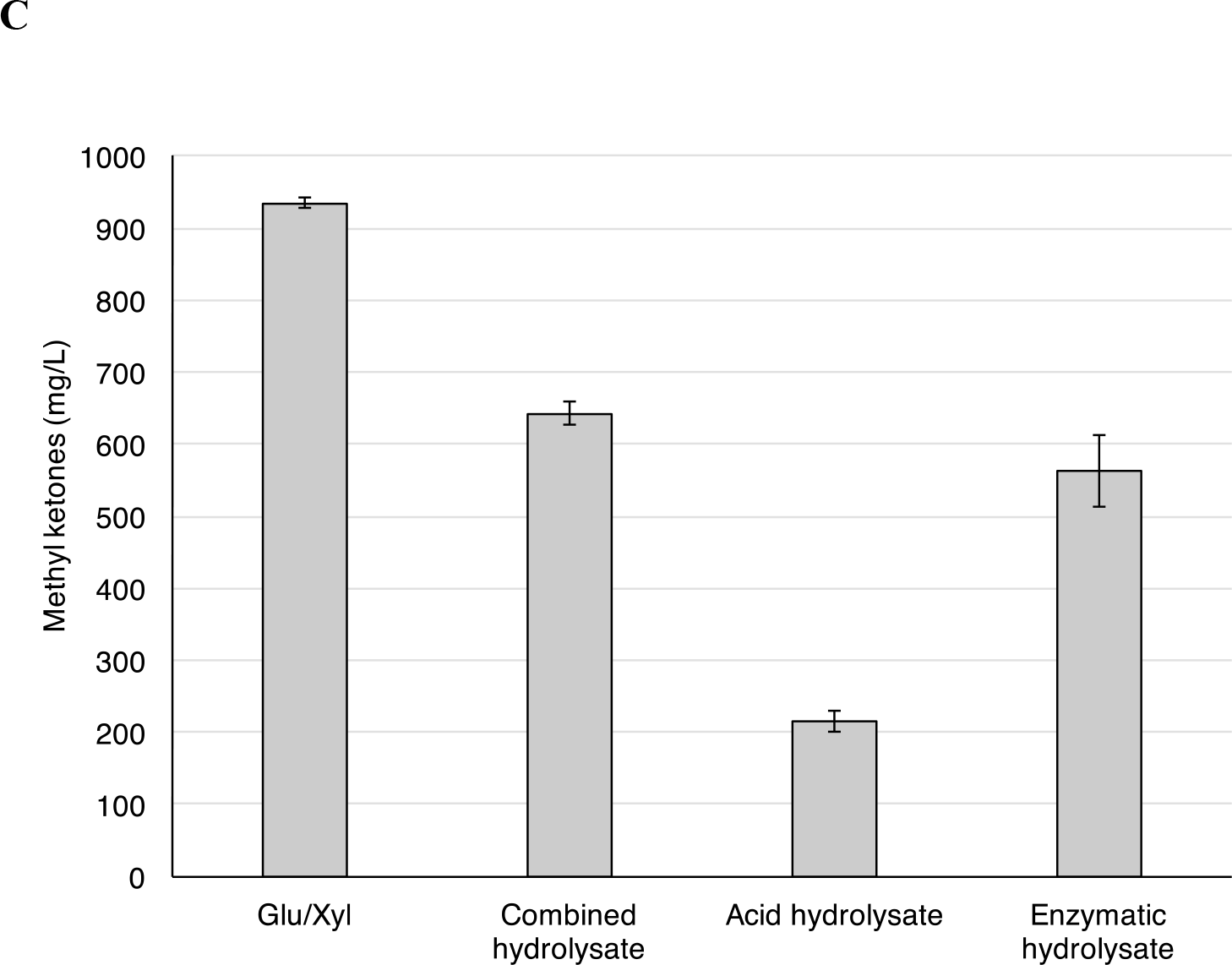
A) HPLC measurements of glucose (Glu) and xylose (Xyl) released by sequential and separate dilute acid and enzymatic hydrolysis of switchgrass. Measurements were performed in triplicate and error reported as standard deviation; B) Methyl ketone production by *P. putida* JD4 with switchgrass hydrolysates. Cultures with switchgrass hydrolysates were compared to a control containing 5 g/L of glucose and 4 g/L of xylose (Glu/Xyl). Methyl ketone production at 48 h was performed as described in the Figure 1 legend; C) *E. coli* EGS1895 methyl ketone production at 72 h with the same switchgrass hydrolysates and Glu/Xyl control as the experiment with P. putida JD4.

Methyl ketone production by *P. putida* JD4 was tested in cultures with the combined and separated switchgrass hydrolysates. Growth, as measured by OD_600_, was better in the cultures with the combined hydrolysate compared to the sugar only control (**Figure S3A**). Final ODs were similar in the cultures grown on each of the separated switchgrass hydrolysates, and were ~50% higher than those attained with the sugar-only control. Methyl ketone production from the combined switchgrass hydrolysate was >7-fold higher (710 mg/L versus 92 mg/L) in the combined hydrolysate compared to the sugar control (**Figure 3B**). The separated switchgrass hydrolysates produced substantially lower titer of methyl ketones, with the acid hydrolysate (170 mg/L) producing higher concentrations that the enzymatic hydrolysate (70 mg/L), despite having ~4.5-fold less glucose in the acid hydrolysate. The comparison of *P. putida* methyl ketone production from the combined and separated switchgrass hydrolysates suggested that the high relative methyl ketone titer obtained for the combined hydrolysate arose from synergistic interactions between components of the acid and enzymatic hydrolysate.

Complementary methyl ketone production experiments using the same combined and separated switchgrass hydrolysates were performed with *E. coli* EGS1895. OD measurements indicated that both the combined and acid hydrolysates promoted better growth than the sugar-only control (**Figure S3B**). In contrast, the enzymatic switchgrass hydrolysate supported lower levels of growth than the sugar controls (OD ~60% of the control). As with the *A. thaliana* hydrolysates, *E. coli* EGS1895 produced lower concentrations of methyl ketones from the combined hydrolysate compared to the sugar control. In contrast to *P. putida*, the enzymatic switchgrass hydrolysate yielded more methyl ketone (550 mg/L) compared to the acid hydrolysate (220 mg/L) and the sum was greater that the production achieved with the combined switchgrass hydrolysate (700 mg/L), which suggests that there were negative interactions between the components of the acid and enzymatic hydrolysate during *E. coli* conversion of the methyl ketones (**Figure 3C**).

### Proteomics

Global proteomic analysis *P. putida* JD4 grown either on hydrolysates from biomass of HCHL line and switchgrass or on glucose-rich control medium was performed to identify the determinants of increased production with these different substrates. Three of the four proteins that constitute the methyl ketone production pathway (*E. coli* FadM, FadB and TesA) were among the top 50 proteins present at highest abundance in the *P.putida* JD4 proteome and were 2-3 fold more abundant in the cultures grown on the HCHL and switchgrass hydrolysates (**Figure 4**) (**Table S1**). The abundance of the *M. luteus* acyl-CoA oxidase encoded by *Mlut_11700* was comparable in all the cultures. A native long chain acyl-CoA dehydrogenase (FadE), which catalyzes the same transformation as the acyl-CoA oxidase, was present at higher abundance in the cultures from switchgrass hydrolysate after 24 h and in the cultures from both the HCHL and switchgrass hydrolysates after 48 h. This protein has previously been characterized as a phenylacyl-CoA dehydrogenase, but can also accept C_14_ and C_16_ acyl-CoAs (McMahon & Mayhew, 2007). FadH, a 2,4-dienoyl-CoA reductase, was present at 3-to-6 fold higher levels in the hydrolysate cultures. This protein has not been characterized in *P. putida*, but functions as a reductase for polyunsaturated acyl-CoA molecules in *E. coli* (You, Cosloy, & Schulz, 1989). The high abundance of FadH suggests that the β-oxidation of unsaturated fatty acids is a key step in methyl ketone production. Small but significant increases in abundance were observed for proteins involved in fatty acid biosynthesis (AccA, AccB, FabD, FabV, FabG, FabB), which are consistent with the observation of increased flux through the fatty acid biosynthesis pathway.

**Figure 4.**
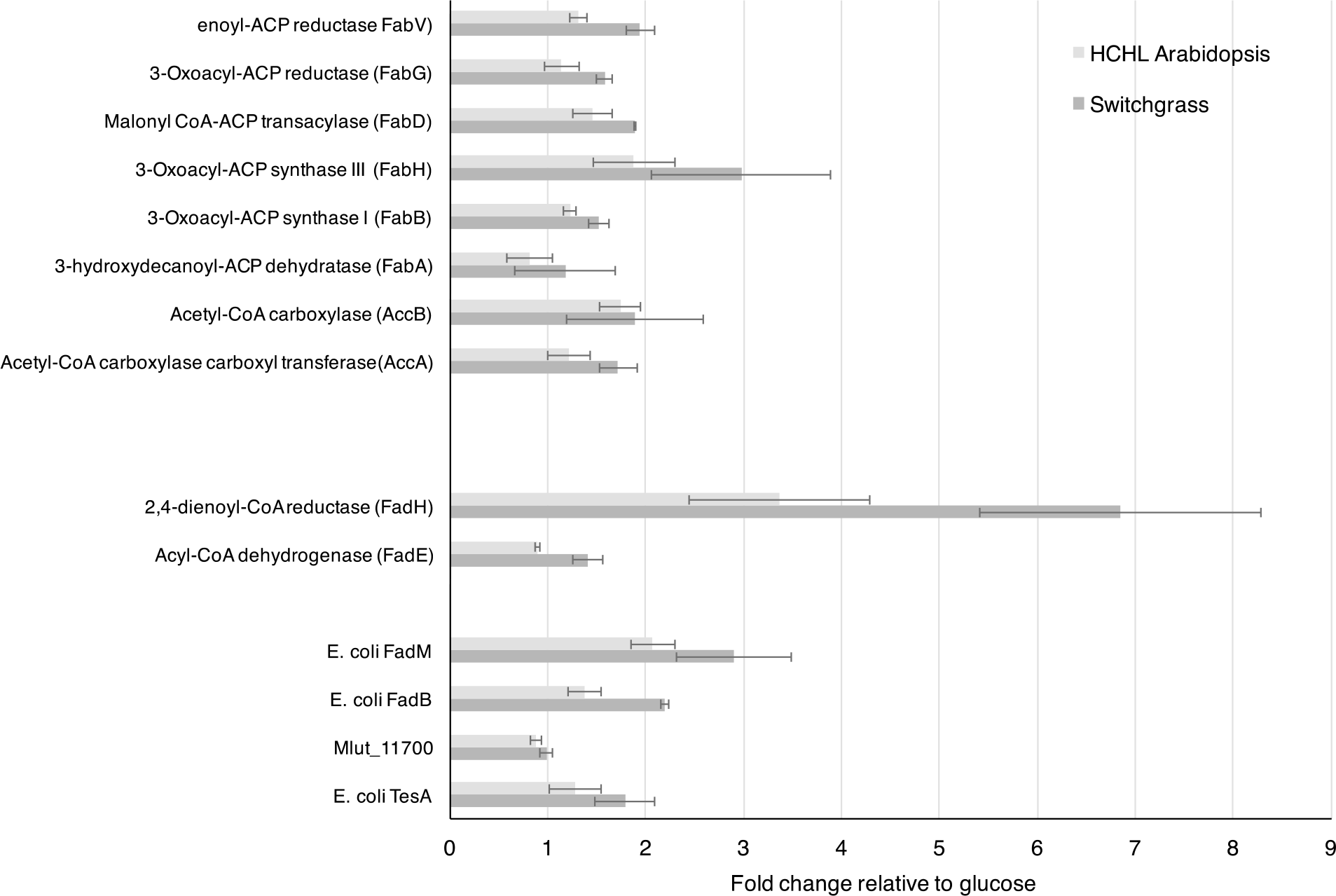
Comparative proteomics of fatty acid metabolism of *P. putida* JD4 during methyl ketone production. Normalized spectral counts of selected proteins (Table S1) were compared for *P. putida* JD4 cultures grown for 24 h with glucose/xylose as carbon sources, or with hydrolysates from *A. thaliana* HCHL and switchgrass biomass. The graph depicts the fold change observed between the cultures grown on hydrolysate and sugar-only control.

Proteins involved in amino acid catabolism were more abundant in the proteomes from the cultures obtained from plant hydrolysate compared to those grown only from glucose (Table S2). Proteins for arginine catabolism: ArcA (arginine deaminase), ArcB (ornithine decarboxylase) and ArcC (carbamate kinase) were present at higher abundances in both cultures from hydrolysate, with the ArcA protein present at ~3 fold higher levels using the *A. thaliana* hydrolysate. For the culture grown on switchgrass hydrolysate, a glutaminase-aspaginase was present at 4-fold higher levels at 24 h compared to the control culture grown from sugar only. In addition, proteins involved in aromatic amino acid catabolism, including: hydroxyphenylpyruvate dioxygenase, homogentisate dioxygenase (HmgA), and fumarylacetoacetate hydrolase (HmgB) were present at higher levels in cultures obtained from the switchgrass hydrolysate.

The proteomic analysis also indicated that other cell wall components besides glucose and xylose were metabolized by *P. putida* in the plant hydrolysates (**Table S3**). Proteins involved in aromatic catabolic pathways were only detected in cultures conducted with the plant hydrolysates. 4-HB hydroxylase, which convert 4-HB to PCA, was present at higher levels in cultures from HCHL hydrolysate, which was consistent with the increased production of 4-HB in the *A. thaliana* HCHL line. Some of the downstream proteins involved in PCA conversion to 3-oxoadipate (PcaH, PcaI) were detected in the proteome, albeit at relatively low abundances. Vanillin dehydrogenase (vdh) and enoyl-CoA hydratase/aldoase, which is involved in ferulate catabolism, were detected at higher abundances in the culture from switchgrass hydrolysates, which reflects the presence of low concentrations of aromatics released by the dilute acid pretreatment and enzymatic hydrolysis (**Table 2**).

**Table 2.** Measured components in biomass hydrolysates.

|  | *A. thaliana*<br>WT (g/L)<sup>a</sup> | *A. thaliana*<br>QsuB (g/L) | *A. thaliana*<br>HCHL (g/L) | Switchgrass<br>(g/L) |
| --- | --- | --- | --- | --- |
| Glucose | 1.94±0.05 | 2.91±0.01 | 2.61±0.02 | 5.02±0.10 |
| Xylose | 2.05±0.05 | 2.41±0.00 | 2.18±0.01 | 4.19±0.10 |
| Arabinose | - | - | 0.044±0.000 | 0.59±0.03 |
| Amino acids<sup>b</sup> | 0.25±0.07 | 0.33±0.00 | 0.85±0.08 | 1.16±0.07 |
| PCA | - | 1.04±0.02 | - | - |
| 4-HB | - | - | 0.20±0.00 | 0.006±0.001 |
| *p*-Coumarate | - | - | - | 0.074±0.002 |
| Ferulate | - | - | - | 0.063±0.002 |
| Acetate | 0.49±0.01 | 0.54±0.00 | 0.53±0.01 | 0.73±0.01 |

<sup>a</sup> This *A. thaliana* WT is the wild-type control of the HCHL line.
<sup>b</sup> This is the total concentration of amino acids. See Figure 4A for more detailed profiles.
“-” indicates the concentration is <0.001 g/L.

### Correlation between amino acid in hydrolysates and methyl ketone production

The elevated levels of amino acid catabolic proteins detected by proteomics for *P. putida* cultivated in the *A. thaliana* HCHL and switchgrass hydrolysates suggested that plant-derived amino acids were critical contributors to the increased methyl ketone titers in plant hydrolysates. Control experiments adding 4-HB to the glucose/xylose control medium did not greatly enhance the methyl ketone production. Inclusion of acetate, present in all the hydrolysates, also did not increase methyl ketone production (**Figure S4**). The concentrations of glucose, the primary substrate for *P. putida*, and arabinose, the inducer for methyl ketone production, were not correlated with methyl ketone production. LC-MS measurements demonstrated that the switchgrass hydrolysate contained the highest concentrations of amino acids (~1.16 g/L) of plant hydrolysates tested for methyl ketone production (**Figure 5A**). The *A. thaliana* HCHL hydrolysate had ~0.85 g/L total amino acids while the *A. thaliana* QsuB hydrolysates and the wild-type hydrolysates had 0.2-0.4 g/L of amino acids (**Table 2**). The amino acid profiles from different biomass shared some common features; the most abundant amino acids in all the hydrolysates were serine, valine, aspartate, phenylalanine and tryptophan. The switchgrass hydrolysate was enriched in aspartate (~0.45 g/L), accounting for the overall increase in amino acid concentration relative to the *A. thaliana* hydrolysates. As validation of this proposed correlation between plant-derived amino acids and methyl ketone production, a sorghum hydrolysate was generated using mild acidic conditions and it contained 0.4 g/L of amino acids with 3.5 g/L of glucose and 2.4 g/l xylose. Incubation of the sorghum hydrolysate with *P. putida* JD3 produced 400 mg/L of methyl ketones, which was consistent with the excellent linear correlation between methyl ketone production and amino acid concentrations for the plant hydrolysates (**Figure 5B**).

**Figure 5.**
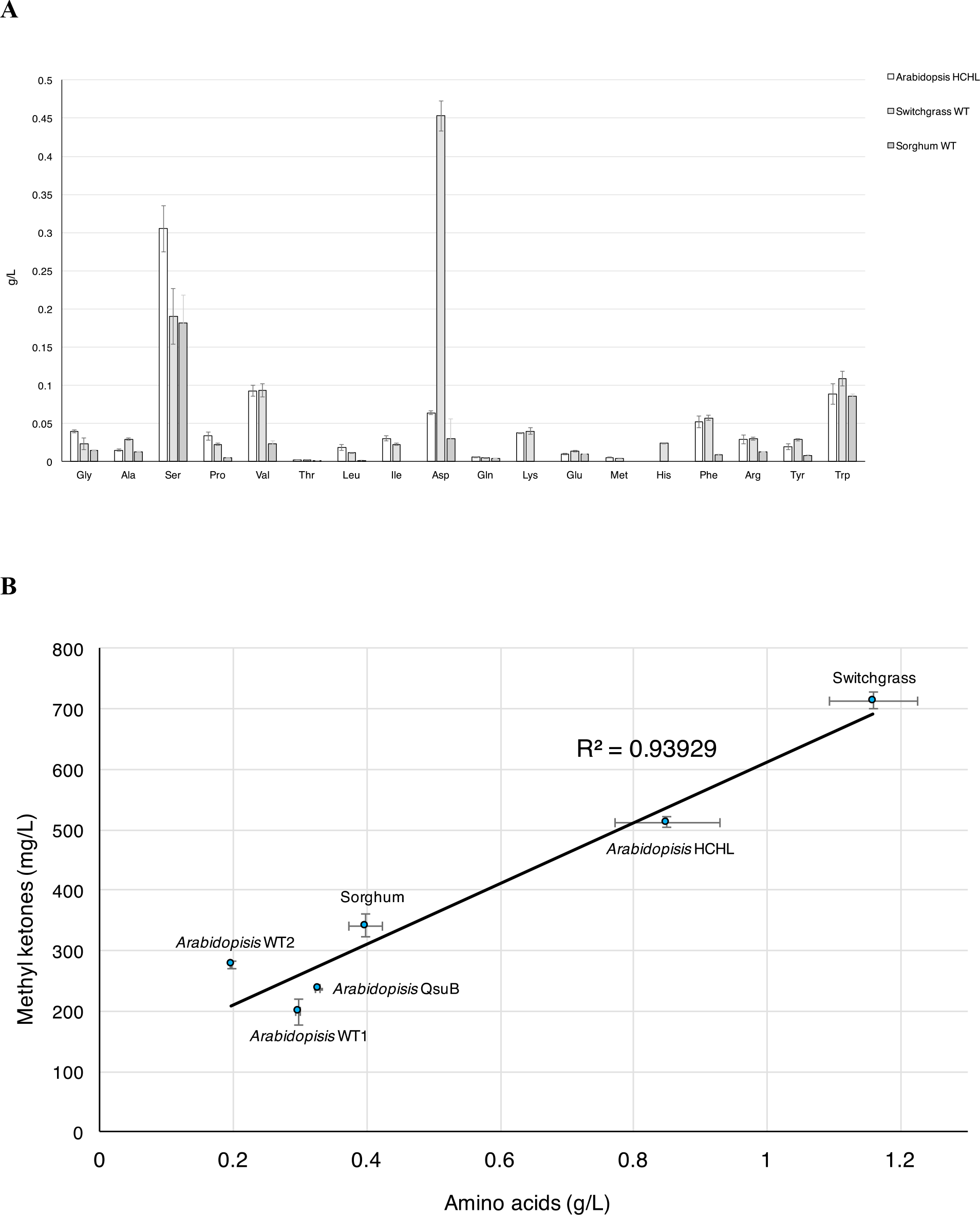
A) LC-MS measurements of amino acid concentrations in acid hydrolysates obtained by pretreatment of *A. thaliana* HCHL line, switchgrass and sorghum. B) Correlation of amino acid concentration in hydrolysates and methyl ketone production by *P. putida* JD4. Methyl ketone production was performed as described in the Figure 1 legend.

### Amino acid-amended medium improves P. putida methyl ketone production

Experiments with plant hydrolysates described above provided evidence that the presence of plant-derived amino acids, produced by acid hydrolysis of the biomass, significantly contributed to the overall increase in methyl ketone production. These results suggested that amending the minimal medium used to produce methyl ketones with amino acids would also increase methyl ketone production. A mixture of the five most abundant amino acids found in the hydrolysates (serine, valine, aspartate, phenylalanine and tryptophan in equal amounts) was added into minimal medium containing glucose (5 g/L). In the absence of glucose, the amino acid mixture was able to support growth and methyl ketone production (193 mg/L at 1.5 g/L amino acids). The addition of glucose resulted in a substantial increase in methyl ketone production (300 mg/L at 0.5 g/L amino acids; 1.1 g/L at 1.5 g/L amino acids) (**Figure 6**), which represented a ~9-fold increase relative to the glucose-only control when MK production from the amino acids was substracted.

**Figure 6.**
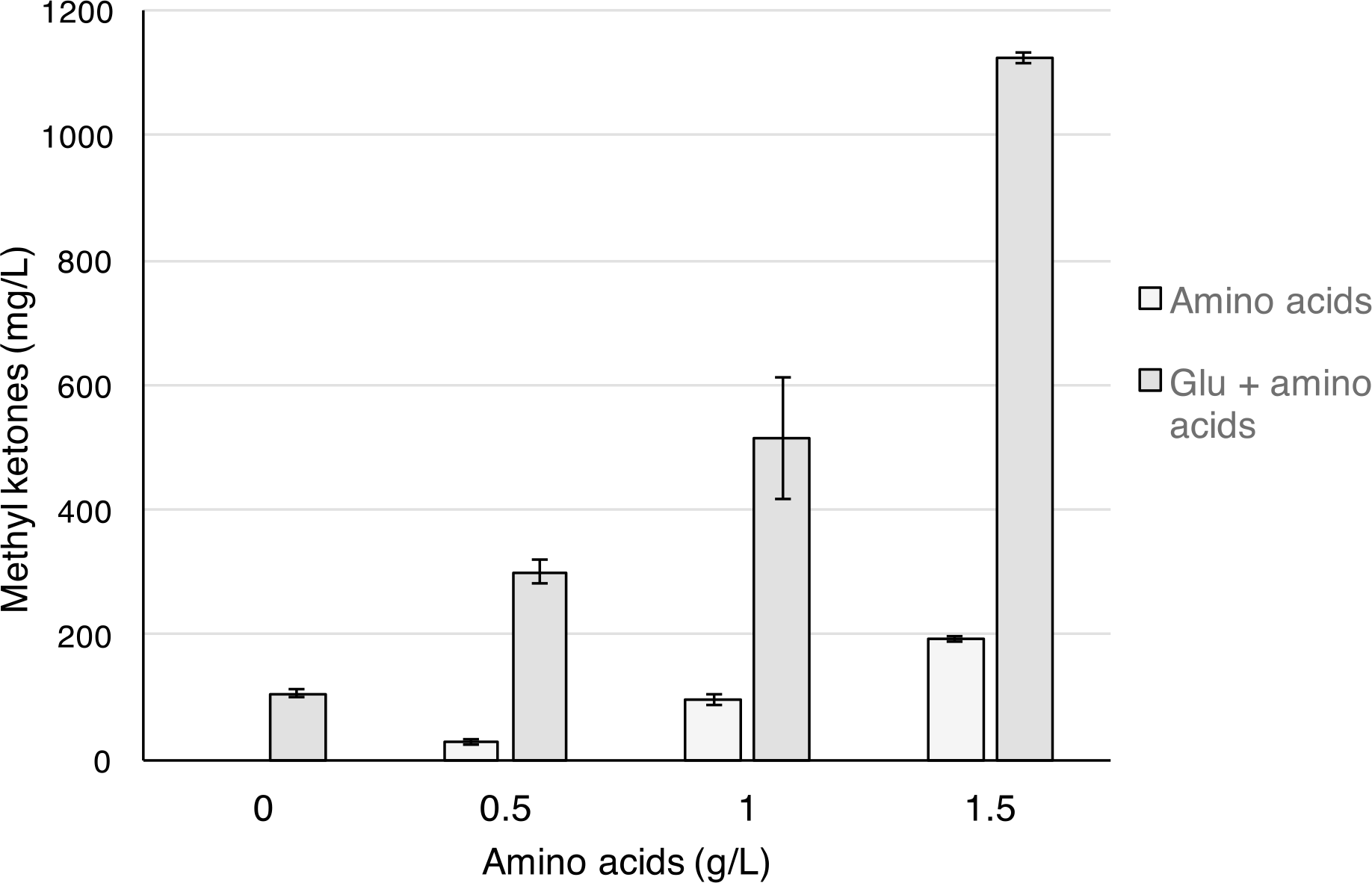
Methyl ketone production in minimal medium supplemented with amino acids. *P. putida* JD4 was cultured on minimal medium with 5 g/L of glucose as the substrate (Glu). The medium was amended with a defined mixture of amino acids (Ser, Val, Asp, Phe and Trp) at total concentrations from 0.5-1.5 g/L. Methyl ketone production at 48 h was performed as described in the Figure 1 legend.

## DISCUSSION

This work was initiated to develop microbial hosts for biofuel and bio-based chemical production that could convert both sugars and aromatics in plant hydrolysates. Here we show that common pretreatment and saccharification protocols (mild acid hydrolysis, combining acid and enzymatic hydrolysates), which generated hydrolysates containing sugars, aromatics, and amino acids, substantially increased production of fatty acid-derived methyl ketones in an engineered *P. putida* strain.

Engineered *P. putida* mt-2 strains produced C_13_ and C_15_ methyl ketones when grown on both glucose and aromatic substrates. The methyl ketone products were highly enriched in unsaturated methyl ketones (>60% C_13:1_ and C_15:1_). In *E. coli*, the C13:0 chain was the most abundant methyl ketone, but there was a substantial proportion of C_13:1_ and C_15:1_ ketones (Goh et al., 2012). In contrast, *R. eutropha* produced C_13:0_ and C_15:0_ at >90% of the total methyl ketone fraction (Müller et al., 2013). Interestingly, the most abundant chain in the membrane fatty acid composition of *P. putida* mt-2 is C_16:0_ (Hachicho, Birnbaum, & Heipieper, 2017), which suggests that the pool of acyl-CoA intermediates that is diverted to methyl ketone production differs from the ACP intermediates that are converted to membrane lipids in *P. putida*. The increase in unsaturated methyl ketones may arise because of cellular stress response, as has been observed for fatty acid production by *E. coli* (Lennen et al., 2011). Additionally, the chain length distribution differs from *mcl*-PHAs produced by *P. putida* KT2440, which has the same chromosomal genotype as *P. putida* mt-2 but lacks the TOL plasmid, when the *phaC* genes are not deleted. *P. putida* KT2440 produced *mcl*-PHAs in which hydroxydecanoate monomers predominated (>50%) when grown on glucose or lignin-related aromatics (*p*-coumarate, ferulate) (Linger et al., 2014)The monomer distribution of *mcl*-PHAs in *P. putida* grown on non-fatty acid substrates is highly dependent by PhaG, a transacylase that converts ACP thioesters to acyl-CoAs and is specific for 3-hydroxydecanoate. Further studies on fatty acid biosynthesis and β-oxidation in *P. putida* are needed to establish the basis for the distribution of methyl ketone products and account for their differences compared to membrane fatty acids and *mcl*-PHA monomers.

Pretreatment of lignocellulosic biomass often generates inhibitors that limit growth and lower product titers and rates relative to those obtained using glucose-containing defined media. This loss of productivity has been observed in the production of succinate from corn stover dilute acid hydrolysate by *Actinobacillus succinogenes* (Salvachúa et al., 2016) and fatty alcohol production from ionic liquid pretreated switchgrass by *Saccharomyces cerevisiae* (Espaux et al., 2017). In particular, dilute acid hydrolysates contain a variety of inhibitors, including phenolics, acetate and furfural, that have been shown to inhibit a variety of hosts for biofuel and biochemical production (Larsson et al., 1999). This phenomenon was observed in this work with the *E. coli* strain that had been engineered for high yield methyl ketone production from glucose. This *E. coli* strain (EGS1895), displayed methyl ketone titers with plant hydrolysates that were 30-60% of the titers obtained for sugar-only controls, consistent with the inhibitory effect of these hydrolysates on bioproduct production. This inhibitory effect on *E. coli* was further supported by the observation that combining the switchgrass acid and enzymatic hydrolysates lowered the methyl ketone titer relative to the sum of the individual hydrolysates indicating a negative synergistic effect (~1.2-fold). *P. putida* strains have demonstrated levels of tolerance to xenobiotic compounds and oxidative stress, so would be expected to respond more favorably to acid hydrolysates than *E. coli*. However, the enhancing effect of plant hydrolysates on *P. putida* methyl ketone production, exemplified by the positive synergy (~3-fold) between the switchgrass acid and enzymatic hydrolysates, indicated that unidentified components were contributing to the methyl ketone production.

Mass spectrometry-based proteomics identified amino acid catabolic proteins at higher levels in the two *P. putida* cultures from plant hydrolysates (*A. thaliana* HCHL and switchgrass) that were consistent with increased amino acid catabolism. Proteomics indicated higher protein levels of most of the gene products involved in fatty acid biosynthesis, especially the heterologous proteins in the methyl ketone pathway, which is consistent with a higher flux toward the fatty acid pathway. The correlation between amino acid content in hydrolysates and methyl ketone production, as well as the increase in methyl ketone production in minimal medium supplemented with amino acid strongly supported the assignment of amino acids as the key stimulative components in the hydrolysates.

The effect of amino acids in plant hydrolysates on microbial performance for bioconversion of lignocellulose is underexplored. The growth of an ethanologenic *E. coli* strain on AFEX-pretreated corn stover hydrolysate was dependent on the presence of amino acids in the hydrolysate, and depletion of these amino acids resulted in a transition to stationary phase. This transition was attributed to an increased requirement for ATP production in the absence of exogenous amino acids (Schwalbach et al., 2012). Potential bioenergy crops traditionally used for forage, such as switchgrass, reed canary grass and alfalfa, have high protein content (5-15%) (Dien et al., 2006) and strategies that integrate amino acid and sugar conversion for hydrolysates derived from these crops may increase the overall efficiency of the biomass to biofuels and biochemicals. This strategy provides a complement to integrating sugar and aromatic metabolism for bioconversion that was the impetus for this study. In both scenarios, *P. putida* is a promising host for bioconversion that possesses capabilities lacking in a widely used host such as *E. coli*.

## CONCLUSION

*P. putida* was successfully engineered to produce C13 and C15 methyl ketones from glucose and lignin-related aromatics, 4-HB and PCA. Methyl ketone production by *P. putida* with *A. thaliana* and switchgrass hydrolysates obtained by dilute acid pretreatment led to the identification of plant-derived amino acids, rather than mono-aromatics, as key enhancing components of these hydrolysates. Shotgun proteomics indicated that the amino acids had a specific inductive effect on proteins involved in fatty acid biosynthesis, leading to a 9-fold increase in methyl ketone titer by amending glucose-containing minimal medium with a defined set of amino acids. This work establishes that the unique metabolic capabilities of *P. putida* are suited to produce high levels of fatty acid-derived biofuels and that proteins in plant biomass may be a promising source of amino acids that increase the conversion efficiency of biomass to biofuels and bio-based chemicals.

## Supporting information

## ACKNOWLEDGEMENTS

This work was performed as part of the DOE Joint BioEnergy Institute (http://www.jbei.org) supported by the U.S. Department of Energy, Office of Science, Office of Biological and Environmental Research, through contract DE-AC02-05CH11231 between Lawrence Berkeley National Laboratory and the U.S. Department of Energy. U.S. Government retains and the publisher, by accepting the article for publication, acknowledges that the U.S. Government retains a nonexclusive, paid up, irrevocable, worldwide license to publish or reproduce the published form of this work, or allow others to do so, for U.S. Government purposes.

## REFERENCES

Andreoni, V., Bernasconi, S., Bestetti, P., & Villa, M. (1991). Metabolism of Lignin-Related Compounds by Rhodococcus rhodochrous - Bioconversion of Anisoin. Applied Microbiology and Biotechnology, 36(3), 410–415.

Beckham, G. T., Johnson, C. W., Karp, E. M., Salvachúa, D., & Vardon, D. R. (2016). Opportunities and challenges in biological lignin valorization. Current Opinion in Biotechnology, 42, 40–53. doi:10.1016/j.copbio.2016.02.030

Beller, H. R., Lee, T. S., & Katz, L. (2015). Natural products as biofuels and bio-based chemicals: fatty acids and isoprenoids. Natural Product Reports, 32(10), 1508–1526. doi:10.1039/c5np00068h

Bokinsky, G., Baidoo, E. E. K., Akella, S., Burd, H., Weaver, D., Alonso-Gutierrez, J., … Keasling, J. D. (2013). HipA-triggered growth arrest and β-lactam tolerance in Escherichia coli are mediated by RelA-dependent ppGpp synthesis. Journal of Bacteriology, 195(14), 3173–3182. doi:10.1128/JB.02210-12

Bokinsky, G., Peralta-Yahya, P. P., George, A., Holmes, B. M., Steen, E. J., Dietrich, J., … Keasling, J. D. (2011). Synthesis of three advanced biofuels from ionic liquid-pretreated switchgrass using engineered Escherichia coli. Proceedings of the National Academy of Sciences of the United States of America, 108(50), 19949–19954. doi:10.1073/pnas.1106958108

Bugg, T. D. H., Ahmad, M., Hardiman, E. M., & Singh, R. (2011). The emerging role for bacteria in lignin degradation and bio-product formation. Current Opinion in Biotechnology, 22(3), 394–400. doi:10.1016/j.copbio.2010.10.009

Choi, K.-H., & Schweizer, H. P. (2005). An improved method for rapid generation of unmarked Pseudomonas aeruginosa deletion mutants. BMC Microbiology, 5, 30. doi:10.1186/1471-2180-5-30

Dien, B., Jung, H., Vogel, K., Casler, M., Lamb, J., Iten, L., … Sarath, G. (2006). Chemical composition and response to dilute-acid pretreatment and enzymatic saccharification of alfalfa, reed canarygrass, and switchgrass. Biomass and Bioenergy, 30(10), 880–891. doi:10.1016/j.biombioe.2006.02.004

Espaux, L. d’, Ghosh, A., Runguphan, W., Wehrs, M., Xu, F., Konzock, O., … Keasling, J. D. (2017). Engineering high-level production of fatty alcohols by Saccharomyces cerevisiae from lignocellulosic feedstocks. Metabolic Engineering, 42, 115–125. doi:10.1016/j.ymben.2017.06.004

Eudes, A., George, A., Mukerjee, P., Kim, J. S., Pollet, B., Benke, P. I., … Loqué, D. (2012). Biosynthesis and incorporation of side-chain-truncated lignin monomers to reduce lignin polymerization and enhance saccharification. Plant Biotechnology Journal, 10(5), 609–620. doi:10.1111/j.1467-7652.2012.00692.x

Eudes, A., Sathitsuksanoh, N., Baidoo, E. E. K., George, A., Liang, Y., Yang, F., … Loqué, D. (2015). Expression of a bacterial 3-dehydroshikimate dehydratase reduces lignin content and improves biomass saccharification efficiency. Plant Biotechnology Journal, 13(9), 1241–1250. doi:10.1111/pbi.12310

Fiorentino, G., Ripa, M., & Ulgiati, S. (2017). Chemicals from biomass: technologicalversus environmental feasibility. A review. Biofuels, Bioproducts and Biorefining, 11(1), 195–214. doi:10.1002/bbb.1729

Gerega, K., Kozłowski, H., Kiss, T., Micera, G., Strinna Erre, L., & Cariati, F. (1987). Cupric complexes with 3,4-dihydroxybenzoic acid. Inorganica Chimica Acta, 138(1), 31–34. doi:10.1016/S0020-1693(00)81177-9

Goh, E.-B., Baidoo, E. E. K., Burd, H., Lee, T. S., Keasling, J. D., & Beller, H. R. (2014). Substantial improvements in methyl ketone production in E. coli and insights on the pathway from in vitro studies. Metabolic Engineering, 26, 67–76. doi:10.1016/j.ymben.2014.09.003

Goh, E.-B., Baidoo, E. E. K., Keasling, J. D., & Beller, H. R. (2012). Engineering of bacterial methyl ketone synthesis for biofuels. Applied and Environmental Microbiology, 78(1), 70–80. doi:10.1128/AEM.06785-11

Goh, E.-B., Chen, Y., Petzold, C. J., Keasling, J. D., & Beller, H. R. (2018). Improving methyl ketone production in Escherichia coli by heterologous expression of NADH-dependent FabG. Biotechnology and Bioengineering, 115(5), 1161–1172. doi:10.1002/bit.26558

González Fernández-Niño, S. M., Smith-Moritz, A. M., Chan, L. J. G., Adams, P. D., Heazlewood, J. L., & Petzold, C. J. (2015). Standard flow liquid chromatography for shotgun proteomics in bioenergy research. Frontiers in Bioengineering and Biotechnology, 3, 44. doi:10.3389/fbioe.2015.00044

Hachicho, N., Birnbaum, A., & Heipieper, H. J. (2017). Osmotic stress in colony and planktonic cells of Pseudomonas putida mt-2 revealed significant differences in adaptive response mechanisms. AMB Express, 7(1), 62. doi:10.1186/s13568-017-0371-8

Hanko, E. K. R., Denby, C. M., Sànchez I Nogué, V., Lin, W., Ramirez, K. J., Singer, C. A., … Keasling, J. D. (2018). Engineering β-oxidation in Yarrowia lipolytica for methyl ketone production. Metabolic Engineering, 48, 52–62. doi:10.1016/j.ymben.2018.05.018

Heinaru, E., Viggor, S., Vedler, E., Truu, J., Merimaa, M., & Heinaru, A. (2001). Reversible accumulation of p-hydroxybenzoate and catechol determines the sequential decomposition of phenolic compounds in mixed substrate cultivations in pseudomonads. FEMS Microbiology Ecology, 37(1), 79–89. doi:10.1111/j.1574-6941.2001.tb00855.x

Johnson, C. W., & Beckham, G. T. (2015). Aromatic catabolic pathway selection for optimal production of pyruvate and lactate from lignin. Metabolic Engineering, 28, 240–247. doi:10.1016/j.ymben.2015.01.005

Kamariotaki, M., Karaliota, A., Stabaki, D., Bakas, T., Perlepes, S. P., & Hadjiliadis, N. (1994). Coordination complexes of iron(III) with 3-hydroxy-2(1H)-pyridinone, 2,3-dihydroxybenzoic acid and 3,4-dihydroxybenzoic acid: preparation and characterization in the solid state. Transition Metal Chemistry, 19(2), 241–247. doi:10.1007/BF00161899

Larsson, S., Palmqvist, E., Hahn-Hägerdal, B., Tengborg, C., Stenberg, K., Zacchi, G., & Nilvebrant, N.-O. (1999). The generation of fermentation inhibitors during dilute acid hydrolysis of softwood. Enzyme and Microbial Technology, 24(3-4), 151–159. doi:10.1016/S0141-0229(98)00101-X

Lennen, R. M., Kruziki, M. A., Kumar, K., Zinkel, R. A., Burnum, K. E., Lipton, M. S., … Pfleger, B. F. (2011). Membrane stresses induced by overproduction of free fatty acids in Escherichia coli. Applied and Environmental Microbiology, 77(22), 8114–8128. doi:10.1128/AEM.05421-11

Linger, J. G., Vardon, D. R., Guarnieri, M. T., Karp, E. M., Hunsinger, G. B., Franden, M. A., … Beckham, G. T. (2014). Lignin valorization through integrated biological funneling and chemical catalysis. Proceedings of the National Academy of Sciences of the United States of America, 111(33), 12013–12018. doi:10.1073/pnas.1410657111

McMahon, B., & Mayhew, S. G. (2007). Identification and properties of an inducible phenylacyl-CoA dehydrogenase in Pseudomonas putida KT2440. FEMS Microbiology Letters, 273(1), 50–57. doi:10.1111/j.1574-6968.2007.00780.x

Meijnen, J.-P., de Winde, J. H., & Ruijssenaars, H. J. (2008). Engineering Pseudomonas putida S12 for efficient utilization of D-xylose and L-arabinose. Applied and Environmental Microbiology, 74(16), 5031–5037. doi:10.1128/AEM.00924-08

Miyada, C. G., Stoltzfus, L., & Wilcox, G. (1984). Regulation of the araC gene of Escherichia coli: catabolite repression, autoregulation, and effect on araBAD expression. Proceedings of the National Academy of Sciences of the United States of America, 81(13), 4120–4124. doi:10.1073/pnas.81.13.4120

Müller, J., MacEachran, D., Burd, H., Sathitsuksanoh, N., Bi, C., Yeh, Y.-C., … Beller, H. R. (2013). Engineering of Ralstonia eutropha H16 for autotrophic and heterotrophic production of methyl ketones. Applied and Environmental Microbiology, 79(14), 4433–4439. doi:10.1128/AEM.00973-13

Ouyang, S.-P., Liu, Q., Fang, L., & Chen, G.-Q. (2007). Construction of pha-operon-defined knockout mutants of Pseudomonas putida KT2442 and their applications in poly(hydroxyalkanoate) production. Macromolecular Bioscience, 7(2), 227–233. doi:10.1002/mabi.200600187

Ouyang, S.-P., Luo, R. C., Chen, S.-S., Liu, Q., Chung, A., Wu, Q., & Chen, G.-Q. (2007). Production of polyhydroxyalkanoates with high 3-hydroxydodecanoate monomer content by fadB and fadA knockout mutant of Pseudomonas putida KT2442. Biomacromolecules, 8(8), 2504–2511. doi:10.1021/bm0702307

Pandey, M. P., & Kim, C. S. (2011). Lignin depolymerization and conversion: A review of thermochemical methods. Chemical Engineering & Technology, 34(1), 29–41. doi:10.1002/ceat.201000270

Rocha, R. C. S., da Silva, L. F., Taciro, M. K., & Pradella, J. G. C. (2008). Production of poly(3-hydroxybutyrate-co-3-hydroxyvalerate) P(3HB-co-3HV) with a broad range of 3HV content at high yields by Burkholderia sacchari IPT 189. World Journal of Microbiology & Biotechnology, 24(3), 427–431. doi:10.1007/s11274-007-9480-x

Salvachúa, D., Mohagheghi, A., Smith, H., Bradfield, M. F. A., Nicol, W., Black, B. A., … Beckham, G. T. (2016). Succinic acid production on xylose-enriched biorefinery streams by Actinobacillus succinogenes in batch fermentation. Biotechnology for Biofuels, 9, 28. doi:10.1186/s13068-016-0425-1

Schwalbach, M. S., Keating, D. H., Tremaine, M., Marner, W. D., Zhang, Y., Bothfeld, W., … Landick, R. (2012). Complex physiology and compound stress responses during fermentation of alkali-pretreated corn stover hydrolysate by an Escherichia coli ethanologen. Applied and Environmental Microbiology, 78(9), 3442–3457. doi:10.1128/AEM.07329-11

Vardon, D. R., Franden, M. A., Johnson, C. W., Karp, E. M., Guarnieri, M. T., Linger, J. G., … Beckham, G. T. (2015). Adipic acid production from lignin. Energy & Environmental Science, 8(2), 617–628. doi:10.1039/C4EE03230F

Vizcaíno, J. A., Csordas, A., del-Toro, N., Dianes, J. A., Griss, J., Lavidas, I., … Hermjakob, H. (2016). 2016 update of the PRIDE database and its related tools. Nucleic Acids Research, 44(D1), D447–56. doi:10.1093/nar/gkv1145

Wu, W., Dutta, T., Varman, A. M., Eudes, A., Manalansan, B., Loqué, D., & Singh, S. (2017). Lignin Valorization: Two hybrid biochemical routes for the conversion of polymeric lignin into value-added chemicals. Scientific Reports, 7(1), 8420. doi:10.1038/s41598-017-07895-1

You, S. Y., Cosloy, S., & Schulz, H. (1989). Evidence for the essential function of 2,4-dienoyl-coenzyme A reductase in the beta-oxidation of unsaturated fatty acids in vivo. Isolation and characterization of an Escherichia coli mutant with a defective 2,4-dienoyl-coenzyme A reductase. The Journal of Biological Chemistry, 264(28), 16489–16495.

Zakzeski, J., Bruijnincx, P. C. A., Jongerius, A. L., & Weckhuysen, B. M. (2010). The catalytic valorization of lignin for the production of renewable chemicals. Chemical Reviews, 110(6), 3552–3599. doi:10.1021/cr900354u

Zhang, F., Carothers, J. M., & Keasling, J. D. (2012). Design of a dynamic sensor-regulator system for production of chemicals and fuels derived from fatty acids. Nature Biotechnology, 30(4), 354–359. doi:10.1038/nbt.2149

