## Supplementary material for "Methyl ketone production by *Pseudomonas putida* is enhanced by plant-derived amino acids"

1. Supplemental Tables S1-S3
2. Supplemental Figures S1-S4

**SUPPLEMNETAL TABLES**

**Table S1**. Normalized spectral counts for *P. putida* JD4 proteins involved in fatty acid metabolism measured during methyl ketone production from glucose and plant hydrolysates.

| | 24 h | | | 48 h | | |
|---|---|---|---|---|---|---|
| | **Glucose** | ***A. thaliana*** **HCHL** | **Switchgrass** | **Glucose** | ***A. thaliana*** **HCHL** | **Switchgrass** |
| *E. coli* ‘tesA | 45 ±4 | 61 ±3 | 93 ±4 | 50 ±4 | 64 ±1 | 75 ±6 |
| *M. luteus* acyl-CoA oxidase (Mlut_11700) | 99 ±2 | 79 ±7 | 99 ±10 | 98 ±3 | 79 ±7 | 92 ±9 |
| *E. coli* FadB | 211 ±18 | 280 ±14 | 492 ±38 | 232 ±5 | 325 ±8 | 492 ±17 |
| *E. coli* FadM | 52 ±5 | 96 ±3 | 152 ±39 | 59 ±2 | 100 ±7 | 128 ±4 |
| Acyl-CoA dehydrogenase (FadE) | 65 ±0 | 52 ±4 | 88 ±7 | 59 ±4 | 76 ±3 | 125 ±4 |
| 2,4 dienoyl-CoA reductase (FadH) | 6 ±1 | 15 ±4 | 40 ±1 | 10 ±2 | 41 ±2 | 50 ±1 |
| Acetyl CoA carboxylase Carboxyl transferase (AccA) | 17 ±2 | 21 ±3 | 30 ±1 | 18 ±2 | 21 ±2 | 30 ±1 |
| Acetyl-CoA Carboxylase (AccB) | 6 ±0 | 8 ±1 | 11 ±2 | 6 ±1 | 9 ±1 | 9 ±1 |
| 3-hydroxydecanoyl-ACP dehydratase (FabA) | 6 ±1 | 4 ±1 | 8 ±1 | 6 ±1 | 4 ±2 | 3 ±1 |
| 3-Oxoacyl-ACP synthase I (FabB) | 21 ±1 | 25 ±3 | 35 ±3 | 21 ±1 | 28 ±1 | 29 ±2 |
| 3-Oxoacyl-ACP synthase III (FabH) | 3 ±1 | 4 ±1 | 9 ±2 | <2 | 5 ±1 | 5 ±1 |
| Malonyl CoA-ACP transacylase (fabD) | 12 ±1 | 16 ±2 | 25 ±2 | 13 ±1 | 16 ±2 | 17 ±0 |

| | | | | | | |
|---|---|---|---|---|---|---|
| 3-Oxoacyl-ACP reductase (FabG) | 14 ±1 | 16 ±1 | 25 ±1 | 8 ±2 | 13 ±2 | 24 ±1 |
| enoyl-ACP reductase (FabV) | 19 ±2 | 22 ±2 | 36 ±2 | 16 ±2 | 26 ±3 | 36 ±2 |

**Table S2.** Normalized spectral counts for *P. putida* JD4 proteins involved amino acid catabolism measured during methyl ketone production from glucose and plant hydrolysates.

| | 24 h | | | 48 h | | |
|---|---|---|---|---|---|---|
| | **Glucose** | ***A. thaliana* HCHL** | **Switchgrass** | **Glucose** | ***A. thaliana* HCHL** | **Switchgrass** |
| Arginine deiminase (ArcA) | 19 ±2 | 55 ±5 | 45 ±4 | 23 ±1 | 69 ±1 | 55 ±1 |
| Ornithine carbamoyltransferase (Arcb) | 9 ±1 | 27 ±2 | 19 ±3 | 9 ±1 | 38 ±5 | 28 ±1 |
| Carbamate kinase (ArC) | 2 ±0 | 8 ±1 | 7 ±1 | 3 ±1 | 10 ±0 | 10 ±1 |
| Glutaminase-aspariginase | 14 ±3 | 22 ±1 | 70 ±2 | 15 ±1 | 26 ±3 | 41 ±3 |
| Hydroxy-phenylpyruvate dioxygenase | 25 ±3 | 17 ±2 | 56 ±6 | 23 ±1 | 18 ±4 | 49 ±1 |
| Homogentisate dioxygenase (HmgA) | 8 ±1 | <2 | 16 ±3 | 6 ±2 | <2 | 17 ±4 |
| fumarylacetoacetate hydrolase (HmgB) | ND[a] | 10 ±3 | 56 ±7 | 2 ±1 | 39 ±3 | 89 ±6 |

[a] Not detected

**Table S3.** Normalized spectral counts for *P. putida* JD4 proteins involved aromatic catabolism measured during methyl ketone production from glucose and plant hydrolysates.

| | 24 h | | | 48 h | | |
|---|---|---|---|---|---|---|
| | **Glucose** | ***A. thaliana* HCHL** | **Switchgrass** | **Glucose** | ***A. thaliana* HCHL** | **Switchgrass** |
| *p*-hydroxybenzoate hydroxylase (pobA) | ND | 15 ±2 | 9 ±1 | ND | 22 ±1 | 11 ±1 |
| Protocatechuate-3,4-dioxygenase (PcaH) | ND | 9 ±2 | 6 ±1 | ND | 10 ±3 | 7 ±1 |
| 3-oxoadipate-CoA transferase (PcaI) | ND | 4 ±1 | 6 ±1 | ND | 4 ±1 | 3 ±1 |
| Vanillin dehydrogenase (vdh) | ND | <2 | 16 ±1 | ND | <2 | 17 ±2 |
| Enoyl-CoA hydratase/aldolase | ND | 11 ±2 | 30 ±2 | ND | 9 ±1 | 30 ±2 |

### SUPPLEMENTAL FIGURES

**Figure S1**. Methyl ketone production from 2% glucose (Glu) and 1.5% 4-HB by *P. putida* strains JD5 and JD6. Culturing, induction and methyl ketone production measurements were performed as described in Figure 1.

**Figure S2.** Optical density at 600 nm of methyl ketone-producing A) *P. putida* JD4; B) *E. coli* EGS1895 cultures grown on *A. thaliana* hydrolysates as described in the Figure 2 legend.

**Figure S3.** Optical density at 600 nm of methyl ketone-producing A) *P. putida* JD4; B) *E. coli* EGS1895 cultures grown on switchgrass hydrolysates as described in the Figure 2 legend.

**Figure S4.** The methyl ketone production of *P. putida* JD4 by adding 0.2 g/L 4-HB and 0.5 g/L acetate in the minimal medium containing 5g/L glucose (Glu) and 4g/L xylose (Xyl).

**Figure S1**

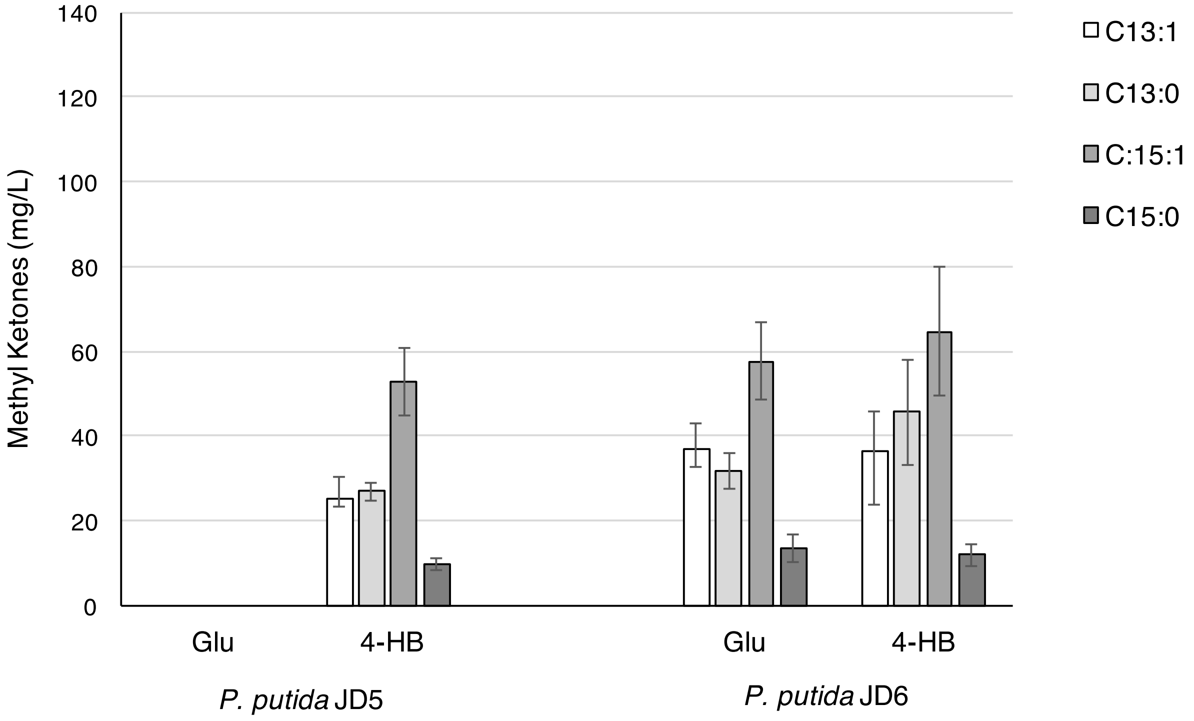

140
120
100
80
60
40
20
0
Methyl Ketones (mg/L)
Glu
4-HB
Glu
4-HB
*P. putida* JD5
*P. putida* JD6
C13:1
C13:0
C:15:1
C15:0

**Figure S2**

**A**

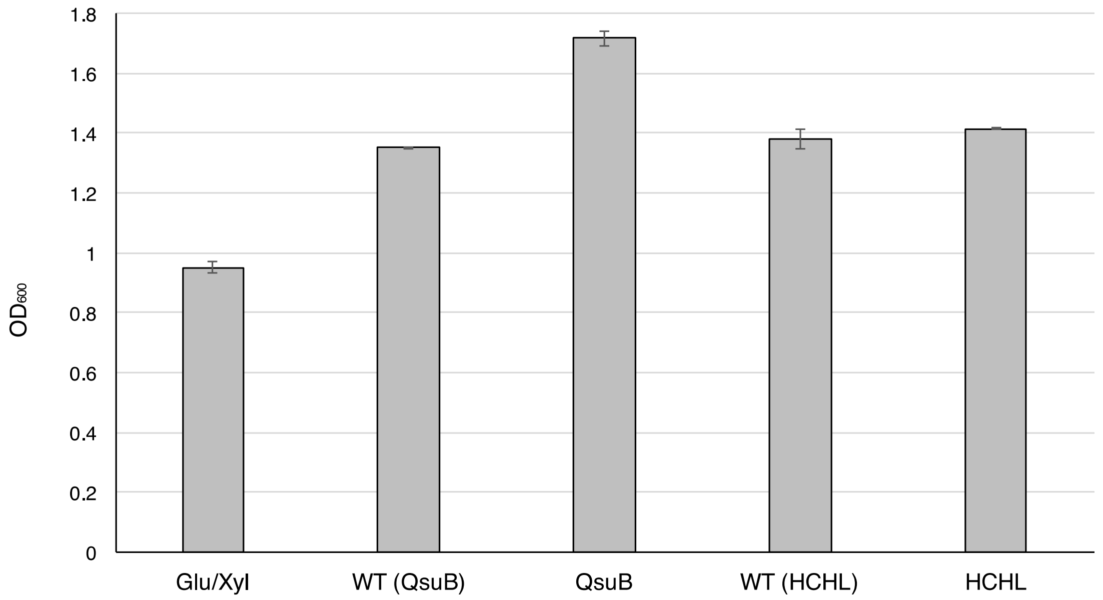
1.8
1.6
1.4
1.2
1
0.8
0.6
0.4
0.2
0
OD600
Glu/Xyl
WT (QsuB)
QsuB
WT (HCHL)
HCHL

**B**

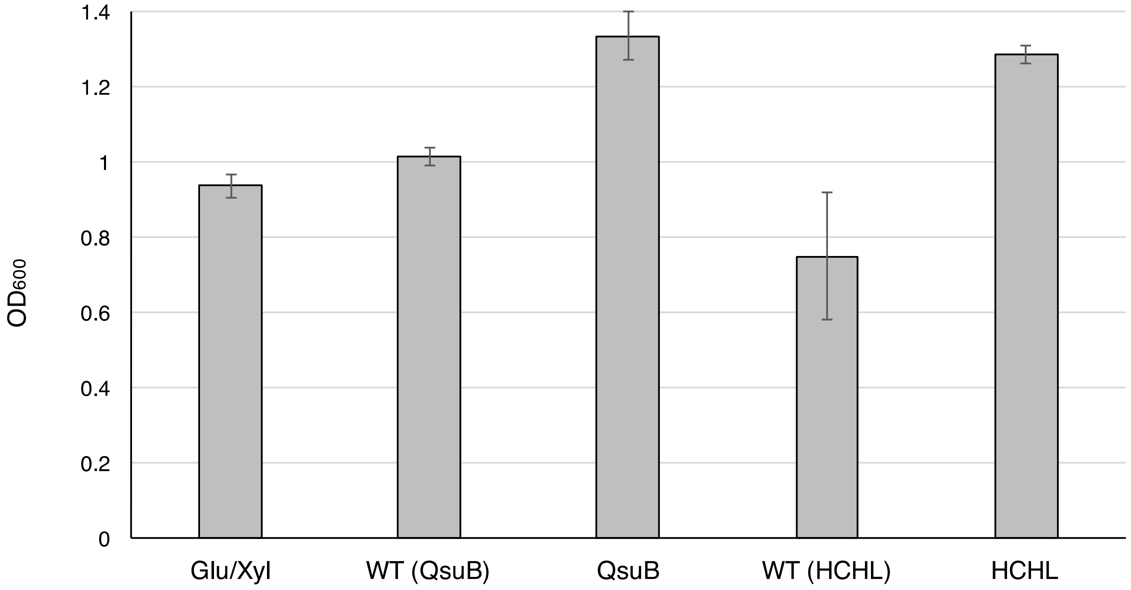
1.4
1.2
1
0.8
0.6
0.4
0.2
0
OD600
Glu/Xyl
WT (QsuB)
QsuB
WT (HCHL)
HCHL

**Figure S3**

**A**

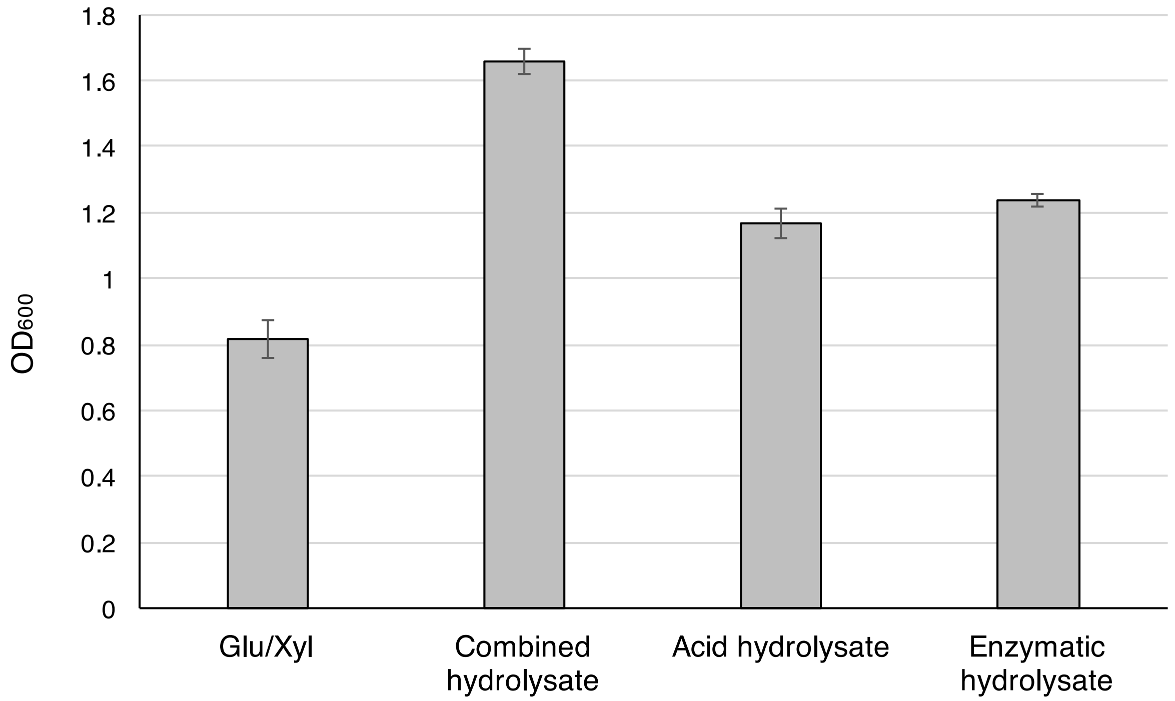

1.8
1.6
1.4
1.2
1
0.8
0.6
0.4
0.2
0
OD600
Glu/Xyl
Combined hydrolysate
Acid hydrolysate
Enzymatic hydrolysate


**B**

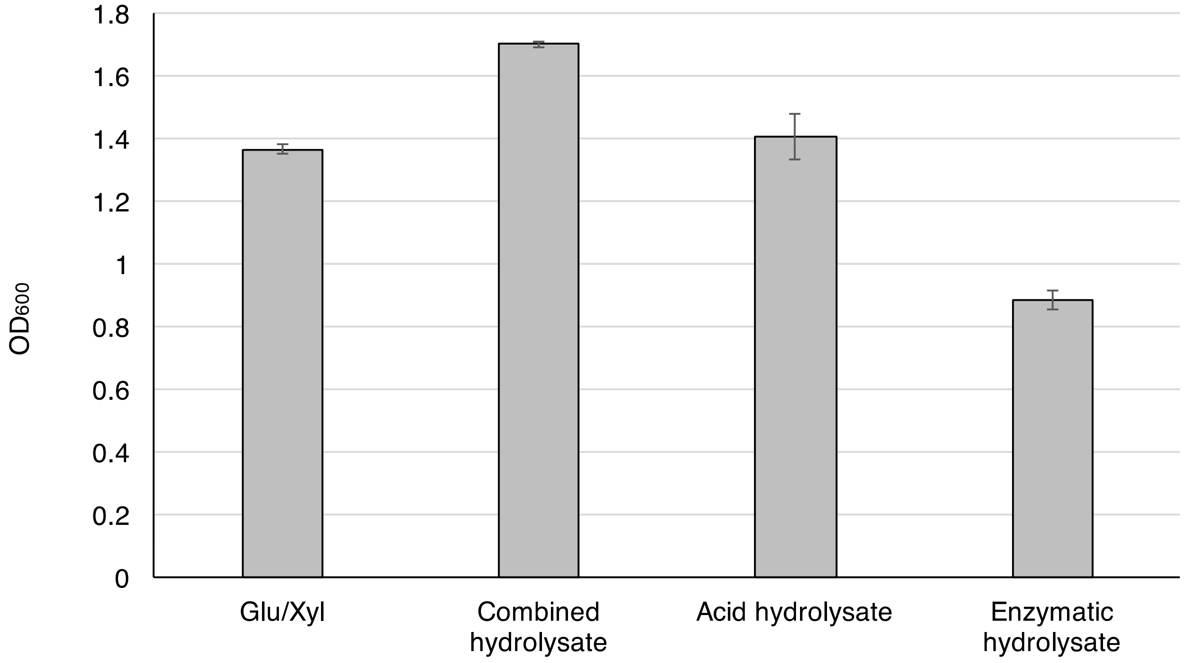

1.8
1.6
1.4
1.2
1
0.8
0.6
0.4
0.2
0
OD600
Glu/Xyl
Combined hydrolysate
Acid hydrolysate
Enzymatic hydrolysate

**Figure S4**

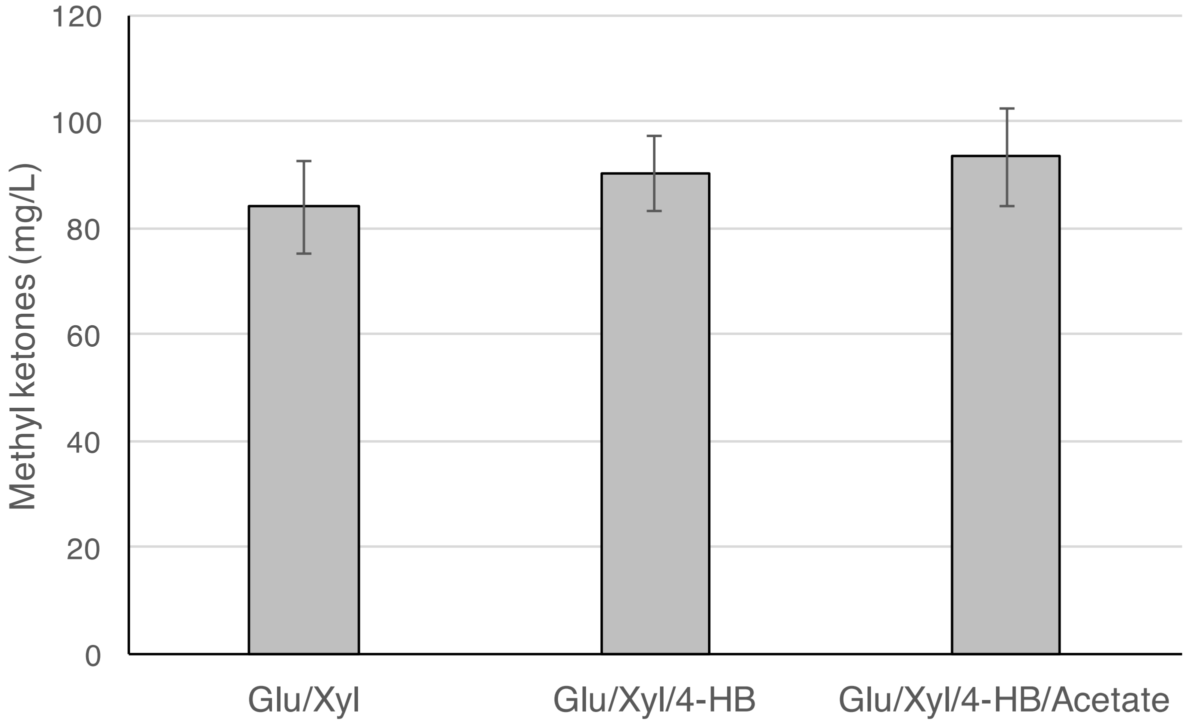
120
100
80
60
40
20
0
Methyl ketones (mg/L)
Glu/Xyl
Glu/Xyl/4-HB
Glu/Xyl/4-HB/Acetate
